# Bait-ER: a Bayesian method to detect targets of selection in Evolve-and-Resequence experiments

**DOI:** 10.1101/2020.12.15.422880

**Authors:** Carolina Barata, Rui Borges, Carolin Kosiol

**Author notes:** These authors contributed equally and will be putting their name first on the citation in their CVs (Carolina Barata, Rui Borges).

## Abstract

For over a decade, experimental evolution has been combined with high-throughput sequencing techniques in so-called Evolve-and-Resequence (E&R) experiments. This allows testing for selection in populations kept in the laboratory under given experimental conditions. However, identifying signatures of adaptation in E&R datasets is far from trivial, and it is still necessary to develop more efficient and statistically sound methods for detecting selection in genome-wide data. Here, we present Bait-ER – a fully Bayesian approach based on the Moran model of allele evolution to estimate selection coefficients from E&R experiments. The model has overlapping generations, a feature that describes several experimental designs found in the literature. We tested our method under several different demographic and experimental conditions to assess its accuracy and precision, and it performs well in most scenarios. Nevertheless, some care must be taken when analysing trajectories where drift largely dominates and starting frequencies are low. We compare our method with other available software and report that ours has generally high accuracy even for trajectories whose complexity goes beyond a classical sweep model.

Furthermore, our approach avoids the computational burden of simulating an empirical null distribution, outperforming available software in terms of computational time and facilitating its use on genome-wide data.

We implemented and released our method in a new open-source software package that can be accessed at https://github.com/mrborges23/Bait-ER.

## 1 Introduction

Natural selection is a complex process that can dramatically alter phenotypes and genotypes over remarkably short timescales. Researchers have successfully tested theoretical predictions and collected evidence for how strong laboratory selection acting on phenotypes can be. However, it is not as straightforward to measure selection acting on the genome. Many confounding factors can lead to spurious results. This is particularly relevant if we are interested in studying how experimental populations adapt to laboratory conditions within tens of generations, in which case we need to take both experiment- and population-related parameters into account.

A powerful approach to gathering data on the genomics of adaptation is to combine experimental evolution, where populations are exposed to a controlled laboratory environment for some number of generations (Kawecki et al., 2012), with genome resequencing throughout the experiment. This approach is referred to as Evolve-and-Resequence (E&R, **fig. 1**). E&R studies are becoming increasingly more common and have already made remarkable discoveries on the genomic architecture of short-term adaptation. Examples of experimental evolution studies include those on yeast (Burke et al., 2014), red flour beetles (Godwin et al., 2017) and fruit flies (Turner et al., 2011; Debelle et al., 2017). The E&R set-up allows for describing the divergence between experimental treatments while accounting for variation among replicate populations (Schlötterer et al., 2015). This is true both at the phenotype and genotype level. Consequently, the optimal approach to finding signatures of selection, is to not only monitor allele frequency changes but to also search for consistent changes across replicates. Moreover, experimental populations are often sampled and pooled for genome sequencing. The motivation for sequencing pooled samples of individuals (pool-seq) is that it is cost-effective and it produces largely accurate estimates of population allele frequencies (Futschik and Schlötterer, 2010). Thus, statistical methods tailored for E&R studies are especially valuable. Notably so when investigating allele frequency trajectories originating from pooled samples of small populations.

**Figure 1:**
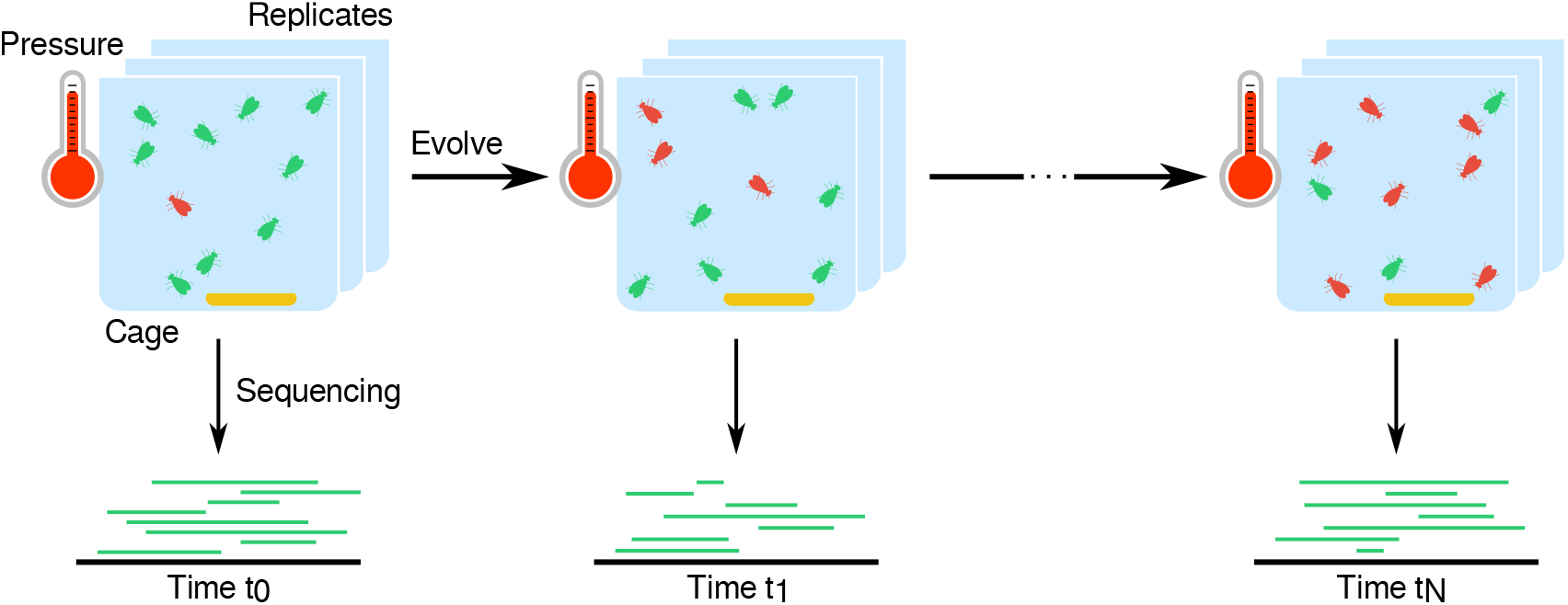
Example of an E&R experimental setup. E&R experiments expose several replicated populations (e.g., of flies, yeast, viruses) to a selective pressure (e.g., temperature, food regimes) for a specific number of generations *t*_*T*_. The replicated populations are surveyed at several time points by whole-genome sequencing, which allows one to quantify changes in allele frequencies over time.

Several statistical approaches have been proposed to analyse these data and detect signatures of selection across the genome. A few such methods consider allele frequency changes between two time points. These simply identify those loci where there is a consistent difference in frequency between time points. One such approach is the widely-used Cochran-Mantel-Haenszel (CMH) test (Cochran, 1954). Such tests are often preferred since they are very fast, which makes them suitable for genome-wide datasets. Other approaches allow for more than two time points: for example, Wiberg et al. (2017) used generalised linear models, and introduced a quasi-binomial distribution for the residual error; and Topa et al. (2015) employed Gaussian Process models in a Bayesian framework to test for selection while accounting for sampling and sequencing noise. While the latter methods use more sophisticated statistical approaches, they remain descriptive with respect to the underlying evolutionary processes. In contrast, mechanistic approaches explicitly model evolutionary forces, such as genetic drift and selection. Such models have the advantage that they can properly account for drift, which may generate allele frequency changes that can easily be mistaken for selection. Indeed, this is usually the case for E&R experimental populations with low effective population sizes (*N*_*e*_), where genetic drift is the main evolutionary force determining the fate of most alleles.

The Wright-Fisher (WF) model is the most used mechanistic model for allele frequencies from time series data. There have been numerous studies that rely on approximations of the WF process, e.g., its diffusion limit (Bollback et al., 2008), a one-step process where there is a finite number of allele frequency states (Malaspinas et al., 2012), a spectral representation of the transition density function (Steinrücken et al., 2014), or a delta method to approximate the mean and variance of the process (Lacerda and Seoighe, 2014). Others have additionally considered the importance of haplotypes arising in a population via mutation (Illingworth and Mustonen, 2012; Nené et al., 2018), or implemented an approximation to the multi-locus WF process over tens of generations (Terhorst et al., 2015). Amongst these methods, most infer selection parameters in the form of selection coefficients, whilst some can also estimate the population size, allele age, mutation rate and even the dominance coefficient. Such parameters are key for understanding the process of genetic adaptation. Nonetheless, there are only a few approaches that couple parameter estimation with explicitly testing for selection (Feder et al., 2014; Terhorst et al., 2015; Iranmehr et al., 2017; Taus et al., 2017; Kojima et al., 2019). While these approaches are useful for detecting selected variants whilst estimating the strength of selection, not all of them are implemented in software packages that can be used genome-wide for E&R experiments.

Most approaches assume linkage equilibrium, and consequently each trajectory is analysed independently from the effects of neighbouring sites. In reality, allele frequencies at linked loci co-vary which causes selection to be overestimated around selected sites. Some have tried to measure the impact of linked selection through analysing autocovariances between adjacent sites (Buffalo and Coop, 2019), and others have investigated the correlation between nearby loci to identify selected haplotypes (Franssen et al., 2017). Whilst these efforts are a step in the right direction, neither approaches directly estimate selection coefficients nor do they test for selection. These two approaches do not rely on modelling evolutionary processes explicitly.

To provide a review of methods that are available for analysing E&R experiments, Vlachos et al. (2019) have produced a comprehensive benchmarking analysis of such methods. Their study compares the programs in terms of overall performance including parameter estimation using simulated data. It features a number of approaches, but not all estimate selection coefficients whilst performing statistical testing for each locus individually. Based on Vlachos et al.’s work, three mechanistic methods are thus particularly relevant in an E&R context: Wright-Fisher Approximate Bayesian Computation (WFABC, Foll et al. (2015)), Composition of Likelihoods for E&R experiments (CLEAR, Iranmehr et al. (2017)) and LLS (Linear Least Squares, Taus et al. (2017)). These methods differ in how they model drift and selection, the inferential approach to estimate selection coefficients, the hypothesis testing strategy, and the extent to which they consider specific experimental conditions (**table 1**). WFABC employs an ABC approach that uses summary statistics to compare simulated and real data. It jointly infers the posterior of both *N*_*e*_ and the selection coefficient at some locus in the genome using allele frequency trajectory simulations. Real and simulated summary statistics must agree to a certain predefined scale. This makes WFABC computationally intensive. CLEAR computes maximum-likelihood estimates of selection parameters using a hidden Markov model tailored for small population sizes. LLS assumes that allele frequencies vary linearly with selection coefficients such that the slope provides the coefficient estimate. Although all three methods have been shown to accurately estimate selection coefficients, they rely heavily on empirical parameter distributions to perform hypothesis testing: (i) WFABC is highly dependent on how accurately the chosen set of summary statistics describes the underlying evolutionary forces determining the observed trajectories; (ii) CLEAR relies on genome-wide simulations to calculate an empirical likelihood-ratio statistic to assess significance; and (iii) LLS computes an empirical distribution of p-values simulated under neutrality. One other common thread amongst these tools is that they do not account for linked selection. Be it background selection or hitchhiking, these software estimate selection without looking into how linked loci might affect other sites’ trajectories. Additionally, the three software vary substantially in computational effort. Therefore, currently available methods are still limited in their use for genome-wide hypothesis testing.

**Table 1:** Currently available software for estimating selection coefficients in E&R experiments.^*a*^.

|  | **WFABC** | **CLEAR** | **LLS** | **Bait-ER** |
| --- | --- | --- | --- | --- |
| Inference approach | Approximate Bayesian computation | Maximum likelihood | Linear least squares* | Bayesian |
| Hypothesis testing | • Bayes factors<br>• Depends heavily on summary statistics | • Likelihood-ratio tests<br>• Empirical p-values based on genome-wide drift simulations | • Empirical simulated p-values based on simulations of allele trajectories | • Bayes factors<br>• Based on the posterior distribution |
| Assumptions | • WF model | • WF model | • WF and Moran model<br>• The allele frequencies vary linearly with the selection coefficients<br>• Weak selection | • Time-continuous Moran model |
| Accounts for replicates | No | Yes | Yes | Yes |
| Accounts for sequencing noise | No | Yes | No | Yes |
| Reference | (Foll et al. 2015) | (Iranmehr et al. 2017) | (Taus et al. 2017) | This study |

<sup>a</sup>The table describes several features of each method namely: i) the approach used for inferring selection coefficients, ii) whether it performs hypothesis testing or not, iii) what sort of assumptions are made about the underlying population genetics model, iv) its overall computational and inference performance, v) whether it accounts for multiple replicate populations, and vi) whether it accounts for sampling variance due to sequencing noise. WF: Wright-Fisher.
\*LLS under the assumption of linearity is equivalent to a maximum likelihood approach.

Here, we propose a new Bayesian inference tool – Bait-ER – to estimate selection coefficients in E&R time series data. It is suitable for large genome-wide polymorphism datasets and particularly useful for small experimental populations. As our new approach was implemented in a Bayesian framework, it gives posterior distributions of any selection parameters while considering sources of experimental uncertainty. Bait-ER jointly tests for selection and estimates selection contrary to other state-of-the-art methods. It does not rely on empirical or simulation-based approaches that might be computationally intensive, and it properly accounts for specific shortcomings of E&R experimental design. As it currently stands, Bait-ER is not concerned with the impact of linked selection. To test Bait-ER and other software, we explore individually simulated trajectories, whole chromosome arm simulations with linkage and an analysis of real data. We show that Bait-ER is faster than other available software, when accounting for hypothesis testing, and still performing accurately in some particularly difficult scenarios.

## 2 Material and Methods

### 2.1 Method outline

E&R experiments produce a remarkable amount of data, namely allele frequencies for thousands to millions of loci. We created a Bayesian framework to infer and test for selection at an individual locus that is based on the Moran model. It estimates the selection coefficient, *σ*, for each allele frequency trajectory, which relies on the assumption that the variant in question is a potential causative locus. The Moran model is especially useful for studies that have overlapping generations, such as insect cage experimental designs (**fig. 1**). Such cage experiments are easier to maintain in the lab and allow for larger experimental population sizes avoiding potential inbreeding depression and crashing populations (Kawecki et al., 2012). Furthermore, Bait-ER combines modelling the evolution of an allele that can be under selection while accounting for sampling noise to do with pooled sequencing and finite sequencing depth. Our method takes allele count data in the widely-used sync format (Kofler et al., 2011) as input. Each locus is described by allele counts per time point and replicate population. The algorithm implemented includes the following key steps:

1. Bait-ER calculates the virtual allele frequency trajectories accounting for *N*_*e*_ that is provided by the user. This step includes a binomial, or beta-binomial, sampling process that corrects for pool-seq-associated sampling noise.
2. The log posterior density of *σ* is calculated for a given grid of *σ*-values. This step requires repeatedly assessing the likelihood function (equation 3 in section 2.2).
3. The log posterior values obtained in the previous step are fitted to a gamma surface (details on surface fitting can be found in **supplementary fig. S1**).
4. Bait-ER returns a set of statistics that describe the posterior distribution of *σ* per locus. In particular, the average *σ* and the log Bayes Factor (BF) are the most important quantities. In this case, BFs test the hypothesis that *σ* is different from 0. Bait-ER also returns the posterior shape and rate parameter values, *α* and *β*, respectively. These can be used to compute other relevant statistics (e.g., credible intervals, variance).

### 2.2 Model description

Let us assume that there is a biallelic locus with two alleles, *A* and *a*. The evolution of allele *A* in time is fully characterised by a frequency trajectory in the state space {*nA*, (*N* − *n*)*a*}, where *n* is the total number of individuals that carry allele *A* (in a population of size *N*). Supposing the allele evolves according to the Moran model, the transition rates for the process are the following

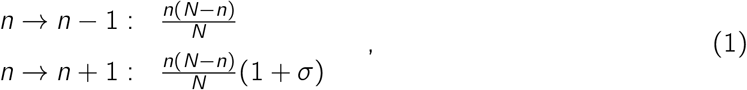

where 1 + *σ* is the fitness of any *A*-type offspring and *σ* the selection coefficient for allele *A*. If *σ* = 0, i.e. *A* is evolving neutrally, then none of the alleles is preferred at reproduction. Let *X*_*t*_ be the number of copies of *A* in a population of *N* individuals; the probability of a given allele trajectory ***X*** can be defined using the Markov property as

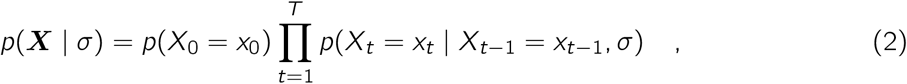

where *T* is the total number of time points measured in generations at which the trajectory was assayed. The conditional probability on the left-hand side of the equation has one calculating 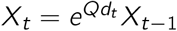, where *Q* is the rate matrix defined in (1) and *d*_*t*_ the difference in number of generations between time point *t* and *t* − 1. The probability of a single allele frequency trajectory can be generalised for *R* replicates by assuming their independence

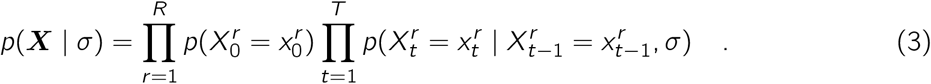

The main caveat for pool-seq data is the fact that it provides estimates for allele frequencies, not true frequencies. For that reason, we assume that the allele counts are generated by a binomial or beta-binomial sampling process which depends on the frequency of allele *A* and the total sequencing depth *C* obtained by pool-seq. We then recalculate the probability of the Moran states given an observed allele count *c*, which becomes the following with binomial sampling

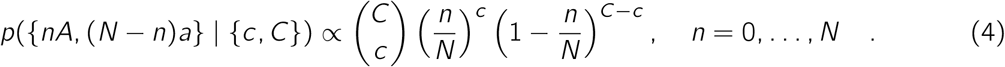

This step is key for it corrects for sampling noise generated during data acquisition. This is particularly relevant for low frequency alleles and poorly covered loci.

### 2.3 Inferential framework

We used a Bayesian framework to estimate *σ*. It requires allele counts and coverage for each time point and replicate population {***c, C***} at each position as input. The posterior distribution can be obtained by

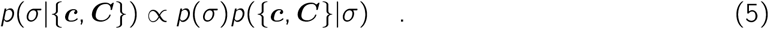

Our algorithm is defined using a gamma prior on *σ*. The posterior cannot be formally obtained, hence we define a grid of *σ* values for which we calculate the posterior density. Estimating the posterior distribution *p*(*σ*|{***c, C***}) is a time consuming part of our algorithm because the likelihood is computationally costly to compute. To avoid this burden, we fit the posterior to a gamma density

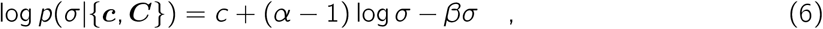

where *α* and *β* are the shape and rate parameters, respectively, and *c* the normalization constant. The gamma fitting represents a good trade-off between complexity, since it only requires two parameters, but its density may take many shapes. As one requires the values of *α* and *β* that best fit the gamma density for further analyses, we find the least squares estimates of *α* and *β* (and *c*), such that the error is minimal. The estimation is as follows

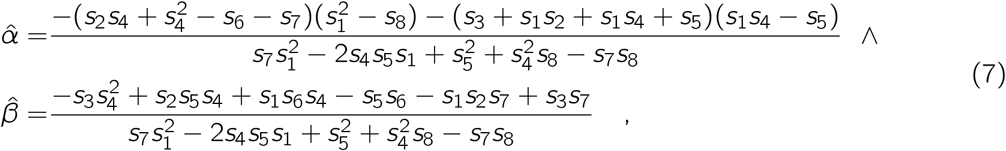

where *s*_1_ = Σ_*i*_ *x*_*i*_ */N, s*_2_ =Σ _*i*_ *y*_*i*_ */N, s*_3_ =Σ _*i*_ *x*_*i*_ *y*_*i*_ */N, s*_4_ =Σ_*i*_ log *x*_*i*_ */N, s*_5_ =Σ _*i*_ *x*_*i*_ log *x*_*i*_ */N, s*_6_ =Σ_*i*_ *y*_*i*_ log *x*_*i*_ */N, s*_7_ =Σ _*i*_ log^2^ *x*_*i*_ */N* and 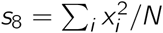. We evaluated the fitting of the gamma density for neutral and selected loci, and observed that a gamma surface with five points describes the log posterior of selected and neutral loci well (**fig. S1**).

Bait-ER was implemented with an allele frequency variance filter that is applied before performing the inferential step of our algorithm. This filtering process excludes any trajectories that do not vary or vary very little throughout the experiment from further analyses. To do that, we assess the trajectories’ frequency increments and exclude loci with frequency variance lower than 0.01. These correspond to cases where trajectories are too flat to perform any statistical inference on. Trajectories such as these typically have both inflated 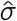 and BFs. This filtering step allows us to improve computational efficiency as we remove trajectories that are statistically uninformative since allele frequencies are essentially constant. Such trajectories are still included in the output file, despite Bait-ER not performing the selection inference step on them. This results in Bait-ER being suitable for large genome-wide datasets without losing any relevant information on trajectories that might be initially flat but can eventually escape drift very quickly.

Bait-ER is implemented in C++ and freely available for download at https://github.com/mrborges23/Bait-ER (accessed on April 9^th^ 2021). Here, we provide a tutorial on how to compile and run Bait-ER, including a toy example with 100 loci taken from Barghi et al. (2019).

### 2.4 Simulated data

We tested our algorithm’s performance under several biologically relevant scenarios using (1) a Moran model allele frequency trajectory simulator, and (2) the individual-based forward simulation software MimicrEE2 (Vlachos and Kofler, 2018).

The Moran model simulator was used, firstly, for benchmarking Bait-ER’s performance across a range of experimental conditions, and, secondly, to compare our estimates of *σ* to those of CLEAR (Iranmehr et al., 2017), LLS (Taus et al., 2017) and WFABC (Foll et al., 2015). We started out by testing Bait-ER under different combinations of experimental and population parameters. A full description of these parameters can be found in **table S2**. Scenarios that explored several experimental designs included those with varying coverage (20x, 60x and 100x), number of replicate populations (2, 5 and 10) and number of sampled time points (2, 5 and 11). In addition to simulating even sampling throughout the experiment, we tested our method on trajectories where we varied sampling towards the start or towards the end of said experiment. Total study length might also affect Bait-ER’s estimation, therefore we tracked allele frequency trajectories for 0.2*N*_*e*_ and 0.4*N*_*e*_ generations.

We set out to compare Bait-ER to other selection estimation software using experimental parameters that resemble realistic E&R designs. Our base experiment replicate populations consist of 300 individuals that were sequenced to 60x coverage. There are five such replicates that were evenly sampled five times throughout the experiment. We then simulated 100 allele frequency trajectories for all starting frequencies and selection coefficients mentioned above. We simulated trajectories for 0.25*N*_*e*_ as well as 0.5*N*_*e*_ generations.

The performance of both CLEAR and LLS was assessed by running the software with a fixed population size of 300 individuals (flag –N=300 and estimateSH(…, Ne = 300), respectively). Additionally, to estimate the selection coefficient under the LLS model, we used the estimateSH(…) function assuming allele codominance (argument h = 0.5). WFABC was tested with a fixed population size of *N*_*e*_ individuals (flag -n 300), lower and upper limit on the selection coefficient of -1 and 1, respectively (flags –min_s -1 and –max_s 1), maximum number of simulations of 10000 (flag –max_sims 10000) and four parallel processes (flag –n_threads 4). The program was run for 1200 seconds, after which the process timed out to prevent it from running indefinitely in case it fails to converge. This caused trajectories with starting allele frequencies of 5% and 1% not to be analysed at all. We have thus only been able to include results for alleles starting at 10% and 50% frequencies.

Finally, we used data simulated with MimicrEE2 (Vlachos and Kofler, 2018) by Vlachos et al. (2019) to benchmark Bait-ER and compare it extensively with other relevant statistical methods. MimicrEE2 allows for whole chromosomes to be simulated under a wide range of parameters mimicking the effects of an E&R set-up on allele frequencies (see **figs. S16 to S21, S25 and S26** for a comparison of population parameters, including nucleotide diversity, with real experimental data). This dataset consisted of 10 replicate experimental populations, and each experimental population consisted of 1,000 diploid organisms evolving for 60 generations. The haplotypes used to find the simulated populations were based on 2L chromosome polymorphism patterns from *Drosophila melanogaster* fly populations (Bastide et al., 2013). Recombination rate variation was based on the *D. melanogaster* recombination landscape (Comeron et al., 2012). 30 segregating loci were randomly picked to be targets of selection. Sites were initially segregating at a frequency between 0.05 and 0.95. Benchmarking Bait-ER using these data also allowed us to look into our method’s robustness when the data generating model is not Moran: the first scenario includes allele frequency trajectories simulated under a Wright-Fisher model of a selective sweep; and the second consists of trajectories simulated under a quantitative trait model with truncating selection. In the former, each of the targets of selection were simulated with a selection coefficient of 0.05. For the latter, 80% of the individuals with the largest trait values were chosen to reproduce.

### 2.5 Application

We applied our algorithm to the recently published dataset from an E&R experiment in 10 replicates of a *Drosophila simulans* population to a hot temperature regime for 60 generations (Barghi et al., 2019). All populations were kept at a census size of 1000 individuals. The experimental regime consisted of light and temperature varying every 12 hours. The temperature was set at either 18°C or 28°C to mimic night and day, respectively. The authors extracted genomic DNA from each replicate population every 10 generations using pool-seq. The polymorphism datasets are available at https://doi.org/10.5061/dryad.rr137kn in sync format. The full dataset consists of more than 5 million SNPs. We subsampled the data such that Bait-ER was tested on 20% of the SNPs. Subsampling was performed randomly across the whole genome.

## 3 Results

### 3.1 Prior fitting with Bait-ER

Bait-ER employs a Bayesian approach outlined in section 2.1 – Method outline – and described in further detail in the section 2.2 – Model description. Bayesian model fitting depends on the prior distribution implemented and requires further testing. Bait-ER uses a gamma prior for which the shape *α* and rate *β* parameters have to be defined beforehand. We tested the impact of uninformative (*α* = *β* = 0.001) and informative (*α* = *β* = 10^5^) gamma priors on the posterior distribution of *σ* under standard (60x coverage, 5 time points and 5 replicates) and sparse (20x coverage, 2 time points and 2 replicates) E&R experiments. Our results show that the prior parameters have virtually no impact on the posterior estimates when *α* = *β* < 100 (**fig. 2** and **supplementary fig. S1**), and thus, by default, Bait-ER sets both prior parameters to 0.001.

**Figure 2:**
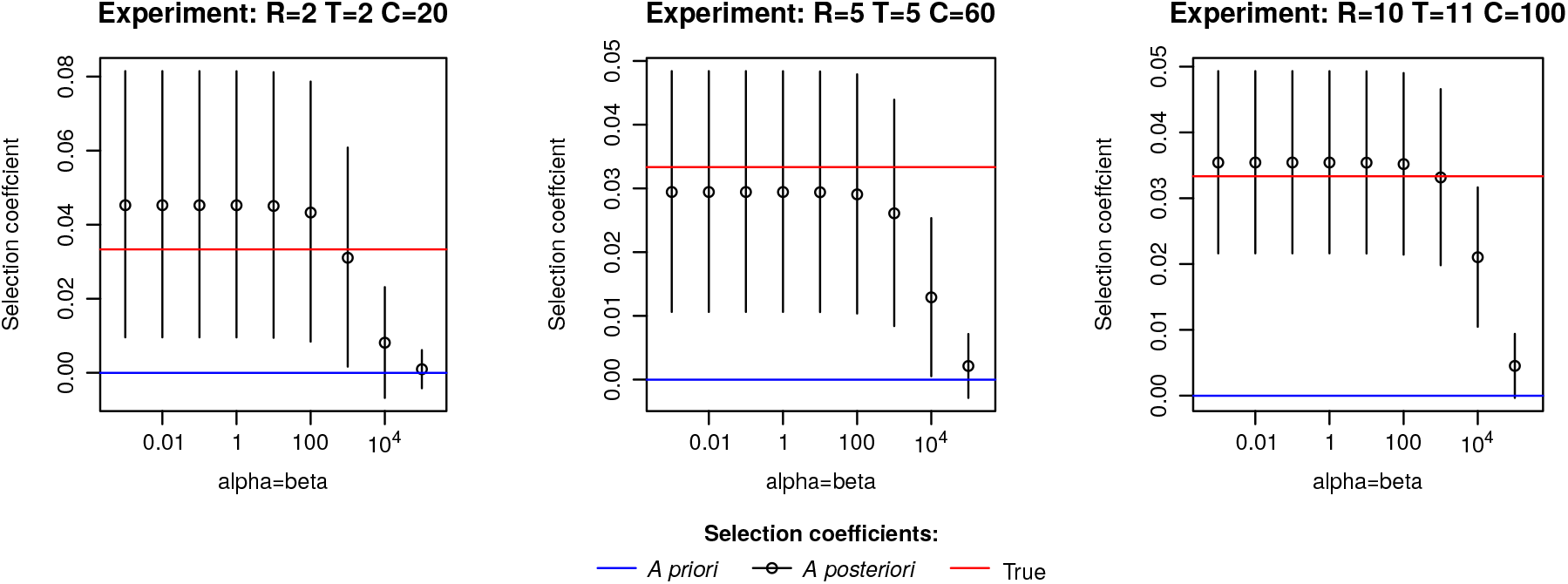
Impact of the prior on the posterior estimates of the selection coefficients. The posterior distribution of *σ* was calculated using gamma priors *G*(*α, β*), where *α* and *β* are the shape and rate parameters. We set *α* = *β* and allowed *β* to vary from 0.001 to 10^5^ (i.e. ranging from a very uninformative to a very informative prior, respectively). The different priors were tested under three E&R experiment scenarios: the first was a sparse experimental design (coverage (C) = 20x, number of time points (T) = 2 and number of replicates (R) = 2), while the second mimicked a standard set up (C = 60x, T = 5 and R = 5). Finally, the third scenario had the most thorough experimental conditions (C = 100x, T = 11 and R = 10). Red lines indicate the true value of *σ*. Blue lines point to the mean of the prior imposed on *σ*. Black lines and points correspond to the posterior mean of *σ* and credibility intervals at 0.95.

Calculating the posterior distribution of *σ* is a computationally intensive step because it requires solving the exponential Moran matrix for several *σ*-values. To reduce the number of times Bait-ER assesses the log-posterior, we fit the posterior density to a gamma distribution. We found that a gamma surface fits the posterior well, and further that five points are enough to provide a good estimate of its surface. This remains valid even for neutral scenarios, where the log-likelihood functions are generally flatter (**fig. S1**).

### 3.2 Impact of E&R experimental design on detecting targets of selection

Bait-ER not only models the evolution of allele frequency trajectories but it also considers aspects of the experimental design specific to E&R studies. Bait-ER can thus be used to gauge the impact of particular experimental conditions in pinpointing targets of selection. We simulated allele frequency trajectories by considering a range of experimental parameters, including the number and span of sampled time points, the number of replicated populations, and coverage.

Each of these settings was tested in different population scenarios that we defined by varying population size, starting allele frequency, and selection coefficient. We assessed the error of the estimated selection coefficients by calculating the absolute bias in relation to the true value. In total, we investigated 576 scenarios (**Supplementary table S2**). Heatmaps in **fig. 3A-C** show the error for each scenario.

**Figure 3:**
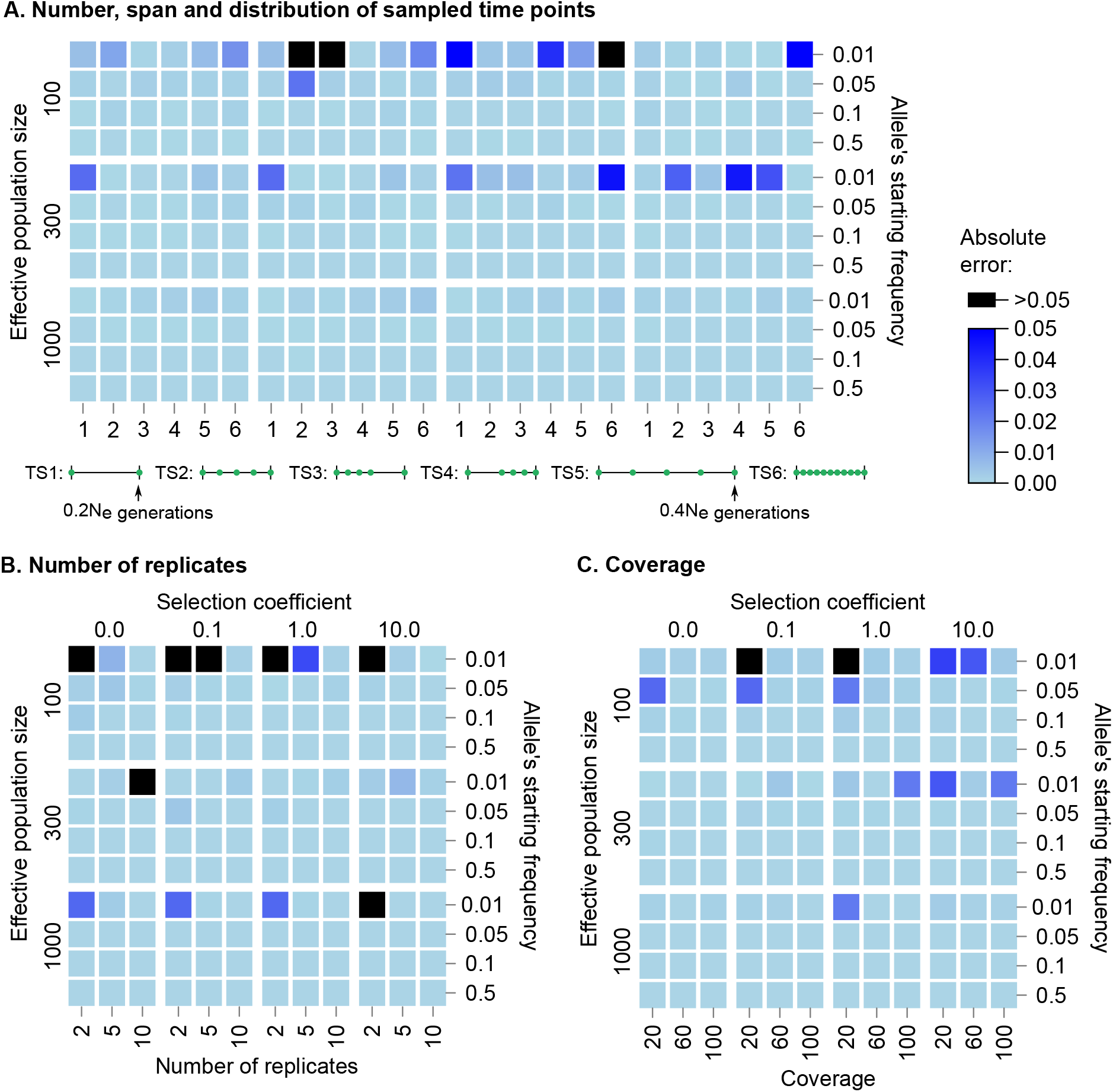
Impact of E&R experimental design on the estimated selection coefficients. Each square of the heatmap represents the error of the estimated selection coefficients, i.e., the absolute difference between the estimated and the true 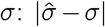, for a range of population dynamics and E&R experimental conditions. **(A)** Number, span and distribution of sampled time points. The six time schemes differ according to the following criteria: most time schemes have five sampling events, except for TS1 and TS6, which have two and eleven time points, respectively; all time schemes have a total span of *N*_*e*_ */*5 generations, except for TS5, which has double the span (2*N*_*e*_ */*5); uniform sampling was used in most scenarios but for TS3, which is more heavily sampled during the first half of the experiment, and TS4, during the second half. The two maximum experiment lengths considered (0.2*N*_*e*_ and 0.4*N*_*e*_) were chosen based on typical E&R experimental designs. **(B)** number of replicates. **(C)** coverage. To test the experimental conditions, we defined a base experiment with five replicates, five uniformly distributed time points (total span of 0.20*Ne* generations) and a coverage of 60x. The complete set of results is shown in **supplementary fig. S2-S5**.

Heatmaps A, B, and C in **fig. 3** show that the initial frequency is a determining factor in the accuracy of 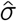 in E&R experiments. We observed that trajectories starting at very low frequencies (around 0.01) may provide unreliable estimates of *σ*. However, 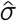’s accuracy on those trajectories can be improved by either increasing the sequencing depth (**fig. S4**) or the number of replicates (**fig. S3**). Similar results have been obtained using other methods such as in Kofler and Schlötterer (2014) and Taus et al. (2017). Designs with high coverage and several replicates may be appropriate when potential selective loci appear at low frequencies (e.g., dilution experiments). Surprisingly, alternative sampling schemes do not seem to substantially impact the accuracy of *σ* (**supplementary text S1**). These results have practical importance because sampling additional time points is time-consuming and significantly increases the cost of E&R experiments.

#### 3.2.1 A note on population size

When using Bait-ER to estimate selection coefficients, one needs to specify the effective population size, *N*_*e*_. However, as effective population size and strength of selection are intertwined, mispecifying *N*_*e*_ will directly affect estimates of selection. The effective population size is often not known at the start of the experiment, but plenty of methods can estimate it from genomic data, e.g., Jonas et al. (2016). To assess the impact of misspecifying *N*_*e*_ on *σ* posterior, we simulated allele frequency trajectories using a fixed population size of 300 individuals. We then ran Bait-ER setting the effective population size to 100 or 1000. By doing so, we are increasing and decreasing, respectively, the strength of genetic drift relative to the true simulated population.

Bait-ER produces highly accurate estimates of *σ* regardless of varying *N*_*e*_ (**fig. 4** and **fig. S5**). Misspecifying it merely rescales time in terms of Moran events rather than changing the relationship between *N*_*e*_ and the number of Moran events in the process. Further, we observed that the BFs are generally higher when the specified *N*_*e*_ is greater than the true value, suggesting that an increased false positive rate. The opposite pattern is observed when the population size one specifies is lower than the real parameter. Additionally, we investigated the relationship between BFs computed with the true *N*_*e*_ and those produced under a misspecified *N*_*e*_. We found that these BFs are highly correlated (Spearman’s correlation coefficients were always higher than 0.99; **fig. 4** and **fig. S5**). Taken together, our results indicate one should use a more stringent BF acceptance threshold if estimates of the real *N*_*e*_ have wide confidence intervals.

**Figure 4:**
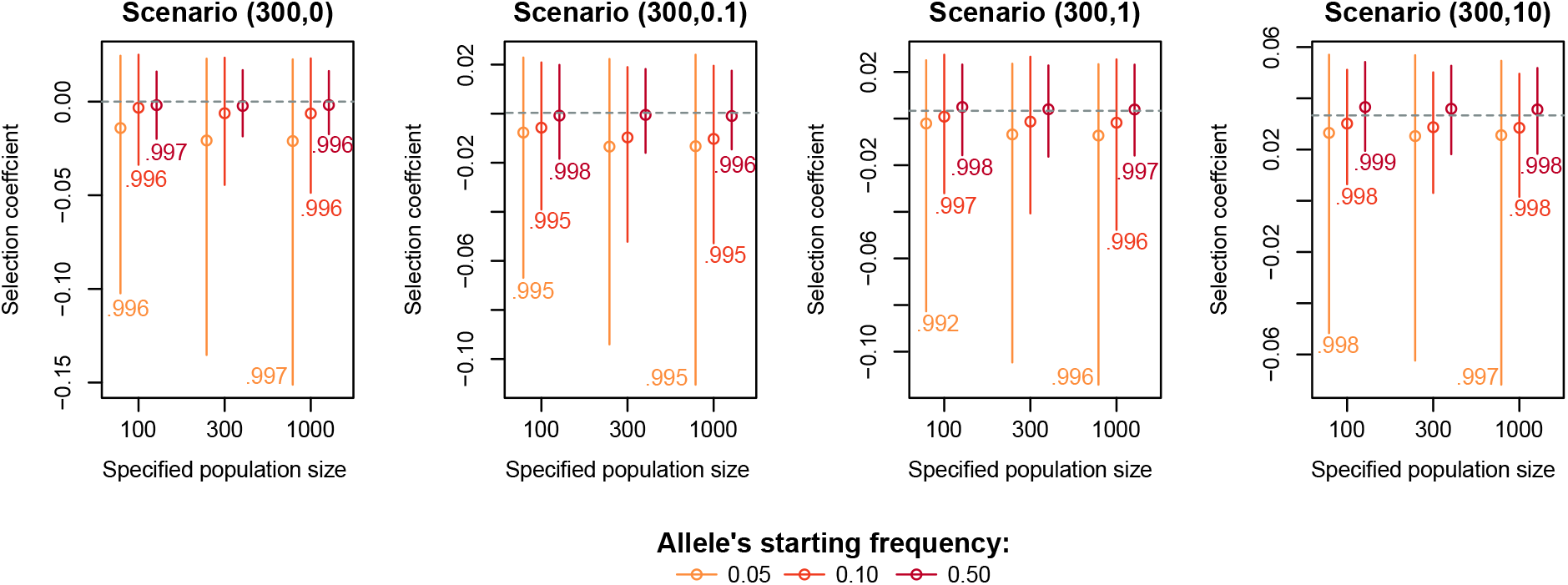
Impact of the user-specified population size on the estimation of selection coefficients. The plots show the distribution of the estimated selection coefficients where the population size is misspecified. Vertical lines and points indicate the interquartile range and median selection coefficient. Each plot represents a specific scenario that was simulated by varying the population size, the true selection coefficient (indicated within brackets (*N*_*e*_, *N*_*e*_ *σ*)) and starting allele frequency (indicated by the yellow-to-red colour gradient). The numbers next to each bar correspond to the Spearman’s correlation coefficient, which correlates the BFs of the 100 replicated trajectories between the cases where we have either under- and overspecified the population size (*N*_*e*_ = 100 or 1000, respectively) and the case where we use the true population size (*N*_*e*_ = 300). Regarding simulated experimental design, we defined a base experiment with five replicates, five uniformly distributed time points (total span of 0.20*Ne* generations) and a coverage of 60x.

Furthermore, we assessed Bait-ER’s computational performance by comparing the relative CPU time while varying several user-defined experimental parameters. We found that increasing *N*_*e*_ affects our software’s computational performance most substantially (31-fold increase in CPU time when increasing the simulated population size from 300 to 1000 individuals; supplementary **table S1**).

### 3.3 Benchmarking Bait-ER with LLS, CLEAR and WFABC

#### 3.3.1 Simulated Moran trajectories

To compare the performance of Bait-ER to that of other relevant software, we set out to simulate Moran frequency trajectories under the base experiment conditions described above. We tested Bait-ER as well as CLEAR (Iranmehr et al., 2017), LLS (Taus et al., 2017) and WFABC (Foll et al., 2015) on 100 trajectories for four starting frequencies (from 1% to 50%) and four selection coefficients (0 ⩽ (*N*_e_*σ* ⩽ 10). All population parameters were tested for both 75 and 150 generations of experimental evolution. **Fig. 5** shows the *σ* estimates for Bait-ER, LLS and CLEAR under two starting frequency scenarios – 10% and 50% – and two *N*_*e*_ *σ*. CLEAR and LLS largely agree with Bait-ER’s estimates of *σ*, even though the level of statistical significance is often not the same. It is evident that LLS produces estimates that are not as accurate as CLEAR’s. This might have to do with the former not explicitly considering sampling bias in pool-seq data as a direct source of error. On the other hand, WFABC systematically disagrees with Bait-ER’s estimates because its distribution is very skewed towards high *N*_*e*_ *σ* (greater than 180; see **fig. S6**). This is perhaps unsurprising given that WFABC does not consider replicate populations nor finite sequencing depth unlike the other three methods. We have included WFABC in our study to compare Bait-ER with another Bayesian method. However, WFABC was not designed for E&R experiments with multiple replicates, hence its poor performance.

**Figure 5:**
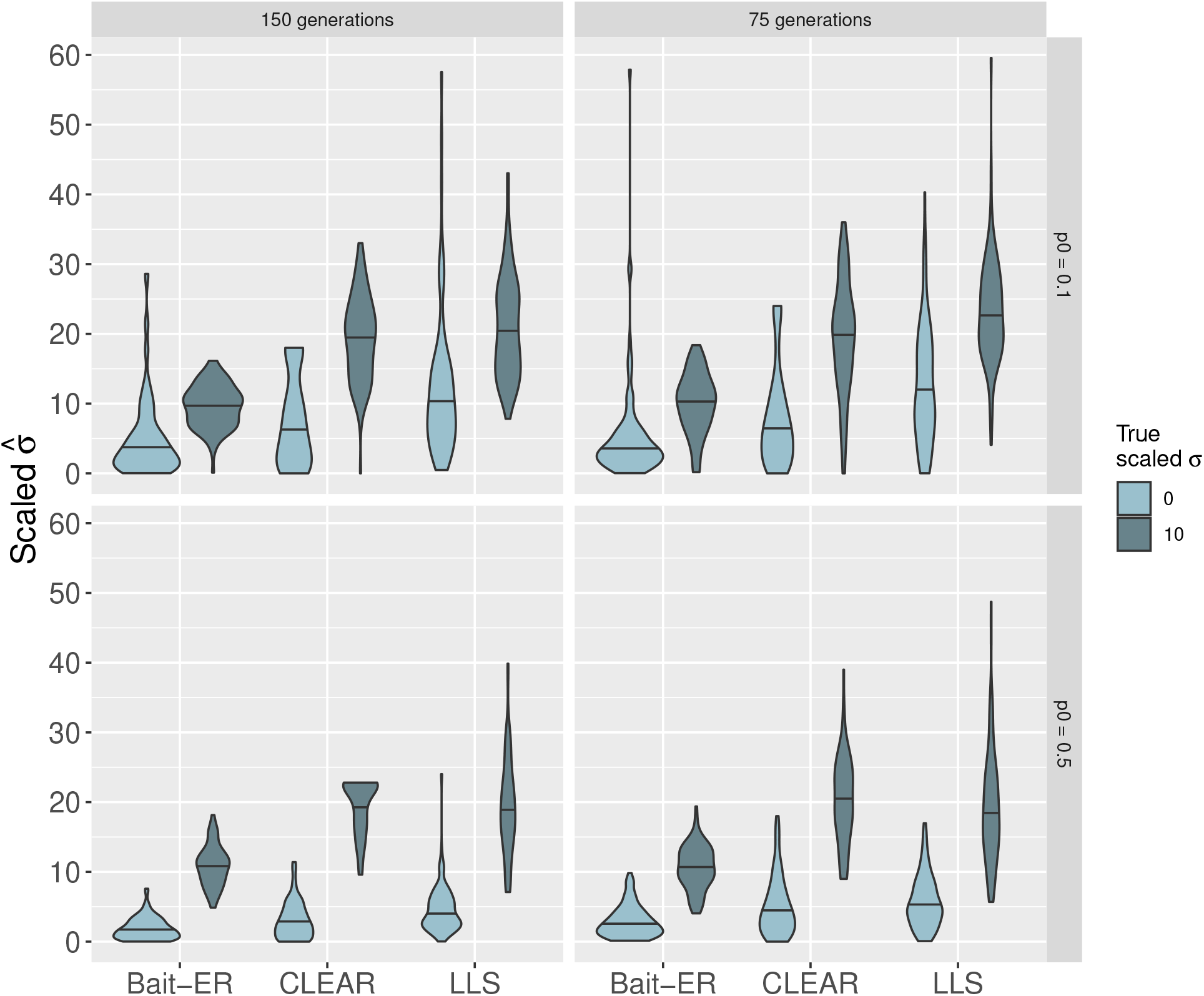
Comparison of estimates of *σ* produced by Bait-ER versus CLEAR and LLS. These plots include estimates for those Moran trajectories simulated with starting frequencies of 10% and 50% (top and bottom row, respectively). Only neutrally evolving (*N*_*e*_*s* = 0) and strongly selected alleles were considered here (*N*_*e*_*s* = 10). The left and right hand side panels correspond to two different experiment lengths: 150 and 75 generations, respectively. LLS returned NA’s for 3 out of 800 trajectories which were excluded from these graphs.

Regarding computational performance, Bait-ER seems to be the fastest of the four methods, even though it is comparable to WFABC (see **fig. 6**). However, we tested WFABC on the first replicate population data rather than the five experimental replicates used for the remaining methods. Additionally, WFABC does not provide any statistical testing output such as a Bayes Factor. In contrast, CLEAR and LLS are slower than the other two approaches. While CLEAR takes less than 40 seconds on average to analyse 100 sites, LLS is the slowest of the four, averaging around 4 minutes. Overall, these results suggest Bait-ER is just as accurate and potentially faster than other currently available approaches, which makes it a good resource for testing and inferring selection from genome-wide polymorphism datasets.

**Figure 6:**
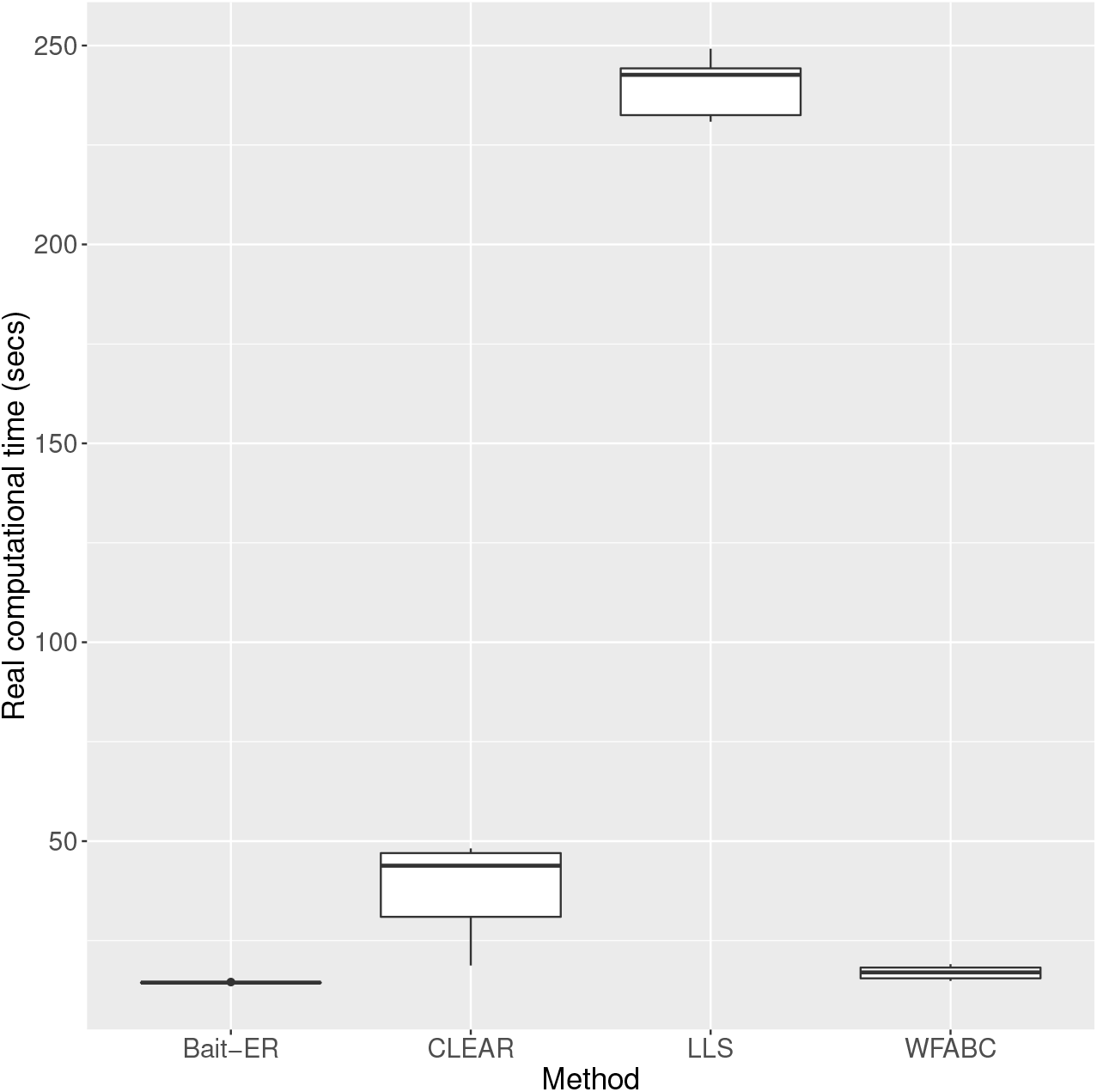
Real computational time for Bait-ER and the other three approaches tested. From left to right, computational time in seconds including both inference and hypothesis testing for Bait-ER, CLEAR, LLS and WFABC is shown here. Similarly to figs. 5 and S6, these boxplots include estimates for those trajectories simulated with starting frequencies of 10% and 50%, as well as both study lengths investigated, i.e. 150 and 75 generations. Four NA’s produced by LLS were again removed from these plots.

#### 3.3.2 Complex simulation scenarios with recombination

For a more comprehensive study of Bait-ER’s performance, we have analysed a complex simulated dataset produced by Vlachos et al. (2019). The authors simulated an E&R experiment inspired by the experimental set-up of Barghi et al. (2019) and used polymorphism data from a *Drosophila melanogaster* population. Vlachos et al. (2019) have produced this dataset to standardise software benchmarking by simulating a series of experimental scenarios that are relevant in an E&R context. We have used it to assess Bait-ER’s performance at inferring selection under linkage and varying recombination rates. In particular, we choose to focus on the classic sweep scenario as well a quantitative trait model with truncating selection, which are two of three complex scenarios simulated in Vlachos et al. (2019). Each experiment had 30 targets of selection randomly distributed along the chromosome arm.

ROC (Receiver Operating Characteristic) curves are compared for five methods, Bait-ER, CLEAR, the CMH test (Agresti, 2003), LLS and WFABC, similarly to fig. 2A in Vlachos et al. (2019). Bait-ER performs well with an average true positive rate of 80% at a 0.2% false positive rate (**fig. 7 (a)**). Its performance is as good as the CMH test’s, but it does underperform slightly in comparison to CLEAR. Bait-ER, CLEAR and the CMH test greatly outperform LLS and WFABC. FIT1 and FIT2 (Feder et al., 2014) are also included for comparison. These methods both use a t-test for allele frequencies and are inaccurate in a classical sweep dataset. A similar picture to that of the sweep simulation emerges for the truncating selection scenario (**fig. 7 (b)**). Bait-ER is amongst the top three methods despite the generating quantitative trait model being completely misspecified during inference. It is only slightly outperformed by CLEAR.

**Figure 7:**
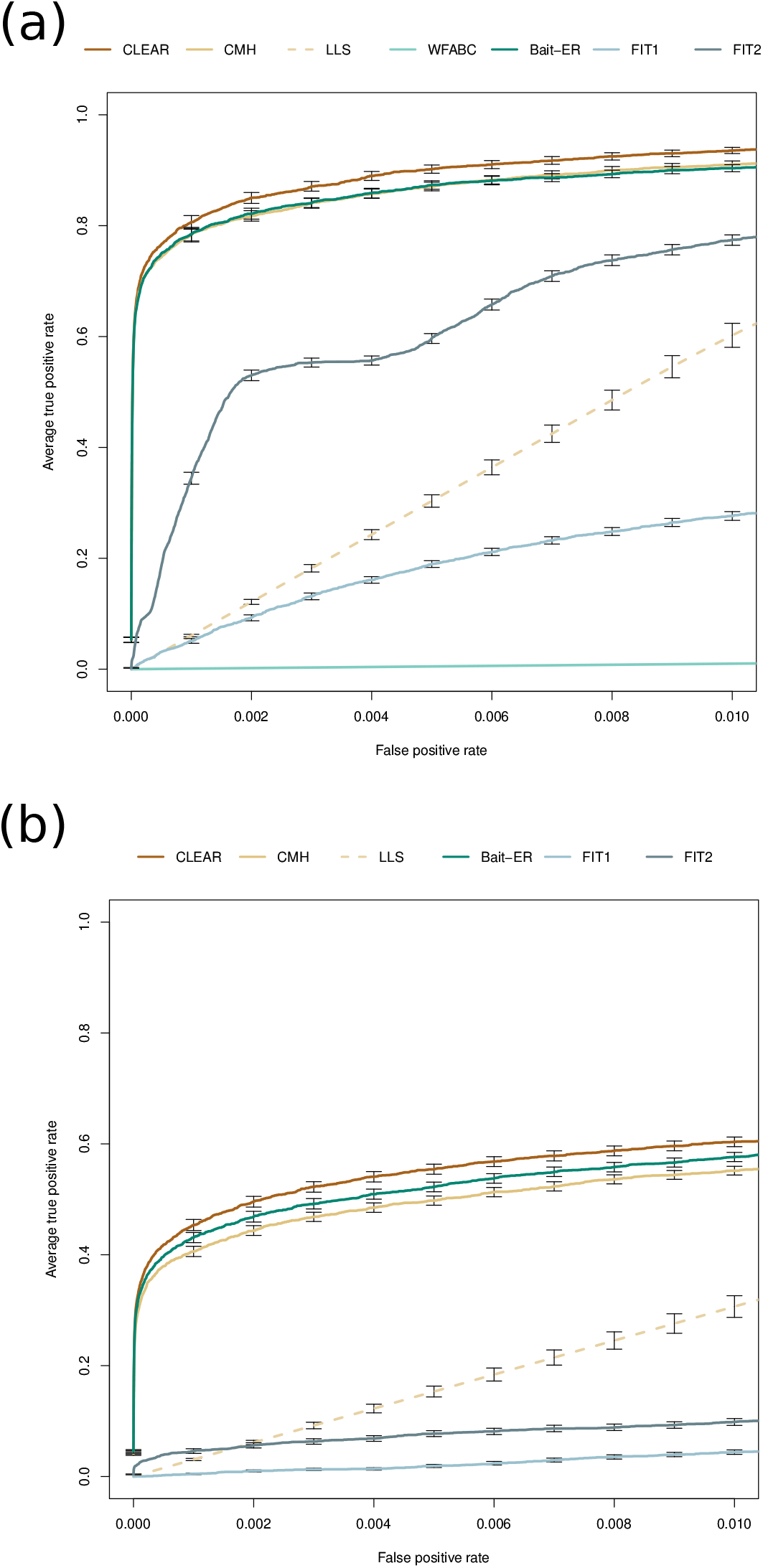
Performance of Bait-ER and other software at testing for selection in data simulated by Vlachos et al. (2019). ROC (Receiver Operating Characteristic) curves for Bait-ER, CLEAR, CMH, LLS, WFABC, FIT1 and FIT2 under (a) the classic sweep scenario and (b) a scenario with truncating selection. Note that LLS and WFABC were run on a subset of SNPs in (a), and that WFABC was not included in (b) for it was prohibitively slow and only finished runs for 29 replicate experiments.

ROC curves serve the purpose of showing how a method’s level of statistical significance compares to other methods’, may it be a p-value or a BF. It addresses whether the method places the true targets of selection amongst its highest scoring hits. While this is informative, it fails to account for the importance of finding a suitable significance threshold. For example, **fig. 7** suggests that Bait-ER and the CHM test perform very similarly. However, the CMH test returns more potential targets than Bait-ER when comparable thresholds are used for both methods (e.g. **fig. 10** that shows the comparison between Bait-ER logBFs and CMH test p-values for a real *D. simulans* dataset). Additionally, whilst the CMH might be more prone to identifying high coverage sites, Bait-ER is not affected by sequencing depth (**fig. S24**). Altogether, this indicates that Bait-ER is more conservative and that the CMH test is more prone to producing false positives.

To assess why Bait-ER seems to be outperformed by CLEAR, we further investigated CLEAR’s selection coefficient estimates. We note that Bait-ER assumes a continuous-time Moran model, whilst CLEAR uses a WF model for inference, much like the simulated data analysed here. Comparison of selection coefficients estimated by Bait-ER and CLEAR showed that Bait-ER is slightly more accurate at estimating true targets’ *σ* (**fig. S7**). In addition, it seems that those trajectories that scored highest with CLEAR are also the highest Bait-ER 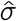 (**fig. S8**). True targets of selection mostly score in the top half of Bait-ER’s *N*_*e*_ *σ* scale (**fig. S23**). Overall, Bait-ER and CLEAR perform to a similar high standard. However, the frequency variance filter implemented in Bait-ER seems to explain our method’s slight underperformance shown in **fig. 7**. Whilst the two method’s false positive rates seem to be comparable, Bait-ER excluded a few selected sites from further analyses as they had changed very little in frequency throughout the experiment. Despite having excluded fewer than 70 (out of 30 targets times 100 experiments) targets of selection, Bait-ER’s filtering step has also classified approximately the same amount of neutral trajectories for being too flat for inferring selection.

Our results also indicate that there is interference between linked selected sites. This phenomenon hinders adaptation as it reduces the fixation probability for each locus - Hill-Robertson Interference, HRI (Hill and Robertson, 1966). It can result in both incomplete and soft sweeps, which are often hard to detect because neither causes the characteristic trough in diversity around causative sites. Bait-ER estimated scaled selection coefficients ranged from 5.85 to 43.2, which suggests each target was under strong selection. Such values should be enough for selection to overcome genetic drift unless there is some degree of interference between selected sites within a 16Mb region. Nevertheless, even with realistic amounts of linked selection, Bait-ER identifies most targets along the chromosome arm and results in narrow peaks of significant BFs (**fig. S15**). For the undetected targets of selection, HRI and inconsistent responses between replicate populations might cause Bait-ER not to perform optimally.

#### 3.4 Analysing E&R data from hot adapted Drosophila simulans populations

We have applied Bait-ER to a real E&R dataset that was published by Barghi et al. (2019). The authors exposed 10 experimental replicates of a *Drosophila simulans* population to a new temperature regime for 60 generations. Each replicate was surveyed using pool-seq every 10 generations. This dataset is particularly suited to demonstrate the relevance of our method, as Barghi et al. (2019) observed a strikingly heterogeneous response across the 10 replicates. The highly polygenic basis of adaptation has proved challenging to measure and summarise thus far. The *D. simulans* genome dataset is composed of six genomic elements: chromosomes 2-4 and chromosome X. For each element, we have estimated selection parameters using Bait-ER (mean 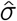 distributions can be found in **fig. S9**). **Fig. 8** shows a Manhattan plot of BFs for the right arm of chromosome 3. We can observe that there are two distinct peaks across the chromosome arm that seem highly significant (BF greater than 9). These two peaks – one at the start and another just before the centre of the chromosome – should correspond to loci that responded strongly to selection in the new lab environment. Such regions display a consistent increase in frequency across replicate populations (see **fig. S22** for the relationship between allele frequency changes and *σ*). Overall, there are only a few other peaks that exhibit very strong evidence for selection across the genome (**fig. S10**). Those are located on chromosomes 2L, 2R and 3L. When compared to the CMH test results as per Barghi et al., Bait-ER’s most prominent peaks seem to largely agree with those produced by the CMH (see **fig. S11**). The same is true for high BF regions on chromosomes 2L and 2R where there are similarly located p-value chimneys at the start of these genomic elements (**fig. S12**). Both Bait-ER and the CMH test did not produce clear signals of selection on chromosomes 3L, 4 and on the X.

**Figure 8:**
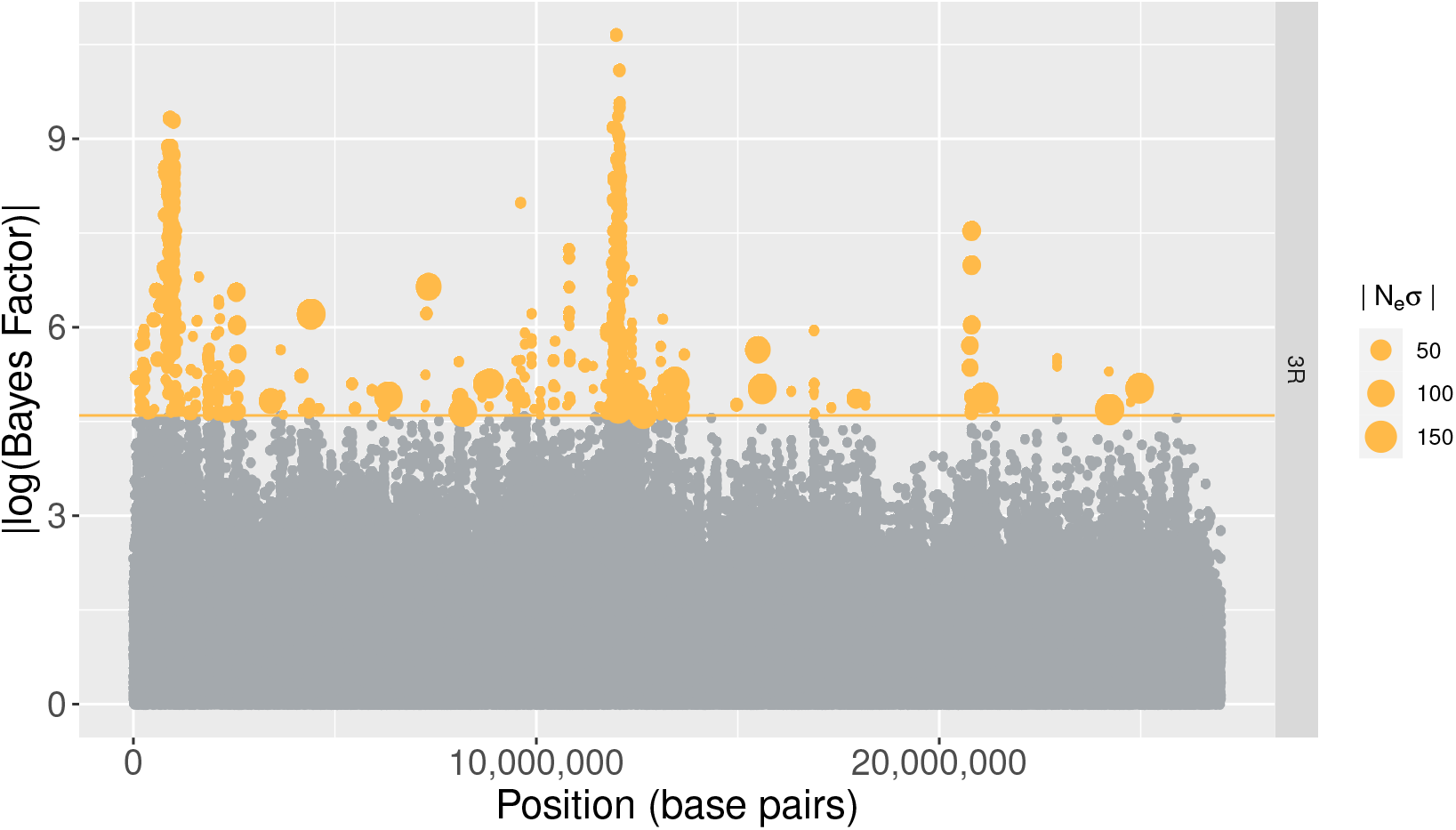
Bayes Factors on chromosome 3R. This Manhattan plot shows log-transformed Bayes Factors computed by Bait-ER for loci along the right arm of the 3^rd^ chromosome in the Barghi et al. (2019) time series dataset. The orange line indicates a conservative threshold of approximately 4.6, which corresponds to log(0.99/0.01), meaning all points in orange have very strong evidence for these to be under selection. The SNPs that are significant at this level are sorted by size according to how strong Bait-ER’s selection coefficients are. In other words, points are sized according to how strong the large selection coefficient is estimated to be.

One of the advantages of Bait-ER is that we have implemented a Bayesian approach for estimating selection parameters, which means we can calculate both the mean and variance of the posterior distributions. To examine both of these statistics, we looked into how the posterior variance varies as a function of mean *σ*. **Fig. 9** shows the relationship between variance and mean selection coefficient for the X chromosome. We observe that the highest mean values also correspond to those with the highest variance. Interestingly, most of those do seem to be statistically significant at a fairly lenient threshold (BF = 2). This suggests that the strongest response to selection, i.e. the highest estimated *σ* values, are also those showing a highly heterogeneous response across replicates. The remaining genomic elements seem to show similar patterns, apart from chromosome 4 (see **fig. S13**). This is consistent with other reports that inferring selection on this chromosome is rather difficult due to its size and low levels of polymorphism (Jensen et al., 2002).

**Figure 9:**
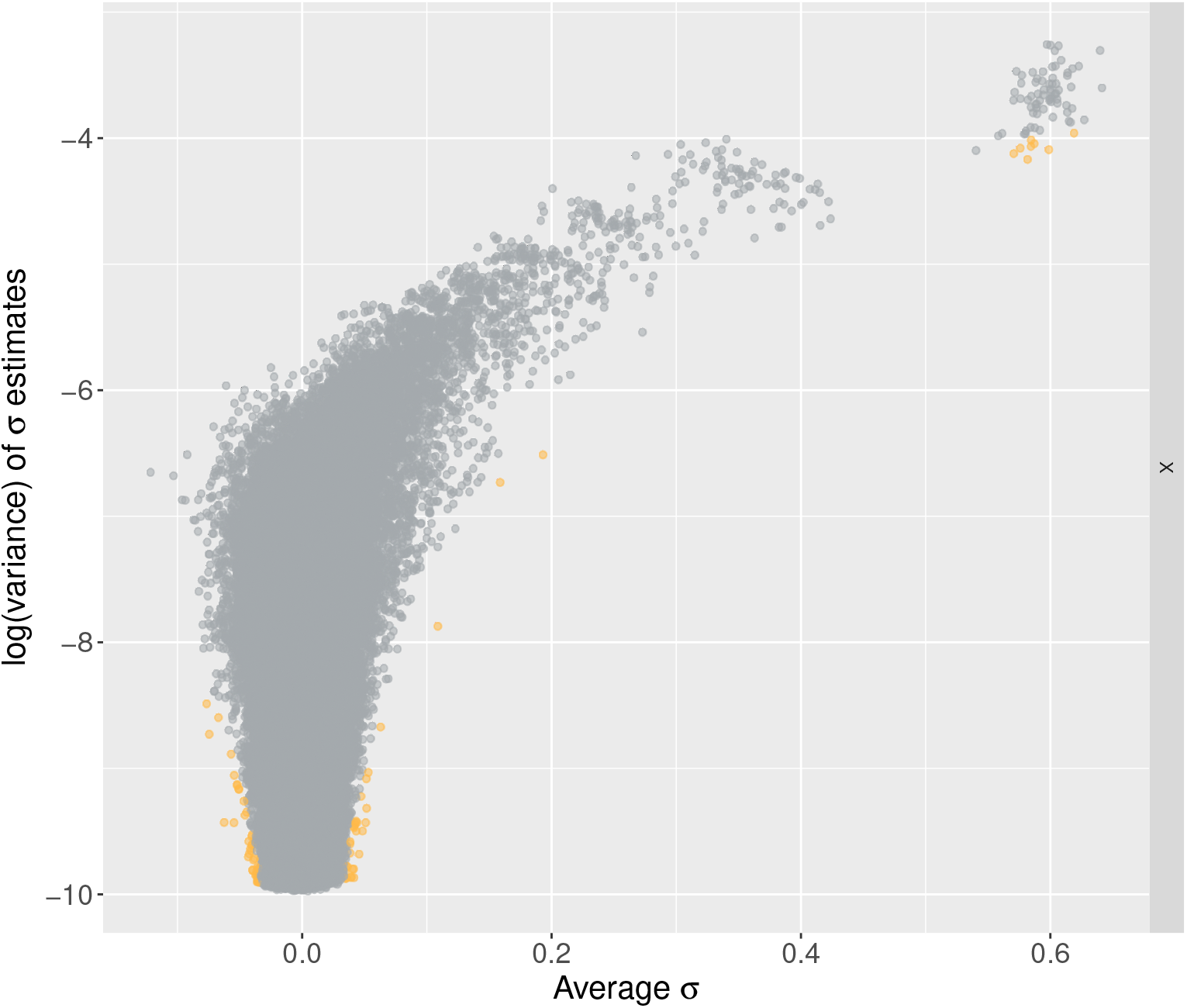
Variance versus mean sigma on the X chromosome. This graph compares log transformed variances in *σ* estimates to average *σ*s. The variance is calculated using the inferred rate and shape parameters for the beta distribution, and the average *σ* is the mean value of the posterior distribution estimated by Bait-ER. Orange coloured points are significant at a conservative BF threshold of log(0.99/0.01), approx. 4.6.

**Figure 10:**
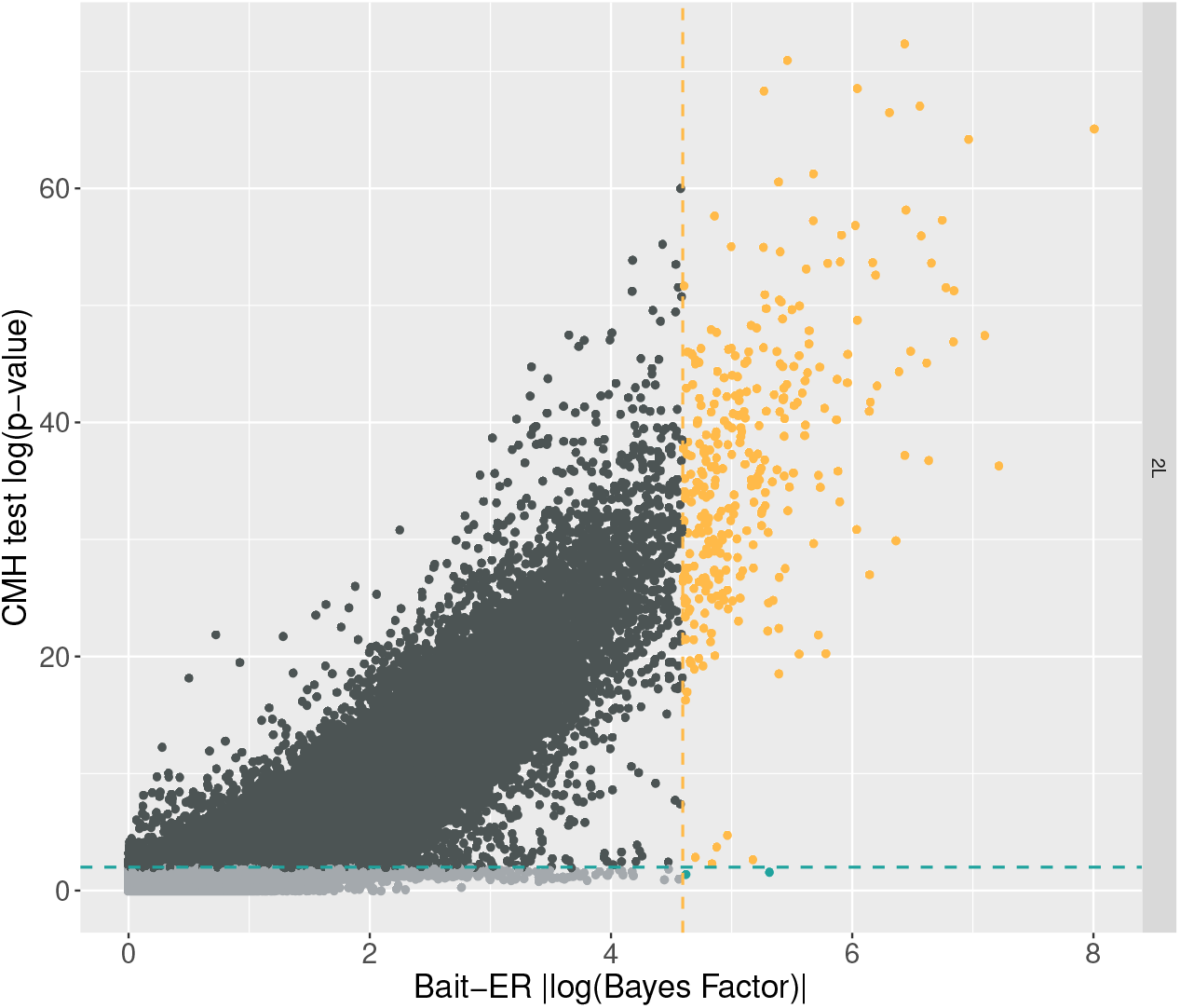
Bait-ER’s Bayes Factors versus CMH test’s p-values on chromosome 2L. Orange coloured points correspond to BFs which are greater than log(0.99/0.01) (approx. 4.6) and p-values less than or equal to 0.01, i.e. those that are considered significant by both tests. Blue coloured points indicated that the computed BF is greater than our threshold, but not significant according to the CMH test. Additionally, dark grey points are significant according to the CMH test, but not to Bait-ER, and light grey points are inferred not significant by both tests.

Finally, we compared the p-values obtained by Barghi et al. (2019) and the BFs computed by Bait-ER. Barghi and colleagues performed genome-wide testing for targets of selection between first and last time points using the CMH test. The tests seem to largely agree for the most significant BFs correspond to the most significant p-values. However, Bait-ER appears to be more conservative than the CMH test. This follows from the finding that there is quite a substantial proportion of loci (less than 10% of all loci) that are deemed significant by a p-value threshold of 0.01, which are not accepted as such by Bait-ER. This is true even for a BF threshold of 2 such as that shown in **fig. 10** for chromosome 2L. Similar patterns are found in other genomic elements (**fig. S14**).

### 3.5 Discussion

One of the main aims of E&R studies is to find targets of selection in genome-wide datasets. For that, we developed an approach that uses time series allele frequency data to test for selection whilst estimating selection parameters for individual loci. As Bait-ER does not rely on simulations for statistical testing, it sets itself apart from other currently available methods. Bait-ER’s implementation of the time-continuous Moran model makes it especially suitable for experimental set-ups with overlapping generations. In addition, we designed Bait-ER to be well suited for small population experiments where genetic drift has a substantial impact on the fate of polymorphisms. This is because random frequency fluctuations can force alleles to be more readily lost and, thus, overlooked by selection. When considering such polymorphisms, our stochastic modelling approach to describe their frequency trajectory is most fitting. We assume that the effect of drift is pervasive and that there is added noise from sampling a pool of individuals from the original population. We show that Bait-ER is faster and just as accurate as other relevant software. Overall, these features make it a desirable approach that can be used in many E&R designs.

Firstly, we addressed Bait-ER’s performance at inferring selection. For that, we carried out a comprehensive analysis of simulated trajectories where we explored the parameter space for coverage, number of experimental replicates, user-defined population size, starting allele frequency and sampling scheme (**figs. S2 to S5**). Our results suggest that Bait-ER’s inference is mostly affected by low starting allele frequencies. This can be overcome should the sequencing depth or the number of experimental replicates be increased. Our simulations show that Bait-ER estimates selection coefficients accurately even if an allele’s starting frequency is low but provided coverage is high and there are at least 5 replicates (**fig. 3**). Although increasing the number of replicates increases the cost of setting up an E&R experiment substantially, improving sequencing depth is certainly within reach. This interesting result might help guide future research. Encouragingly, Bait-ER performed well with small manageable population sizes, suggesting replication is key, but large populations are not necessarily required for achieving good results.

We also assessed Bait-ER’s performance on a complex chromosome arm dataset simulated by Vlachos et al. (2019). We then compared it to other selection inference programs of which most are suited for time series allele frequency data. Despite numerous similarities, they vary substantially in terms of model assumptions and what sort of experimental set-up they are a good fit for. For example, WFABC seems to underperform in comparison with the other methods for E&R experiments. This is likely to be the case because it was modelled for relatively large populations. As Foll et al. (2015) show in their original study, WFABC is accurate for population sizes of 1000 individuals and for both weak and strong selection coefficients. Despite this being low in comparison to experiments in bacteria or yeast, which easily range from 10^5^ to 10^8^, that is not the standard population size we consider in our work. Bait-ER has been shown to perform well for such large populations (see bottom rows of each graph in **fig. 3**), as well as small census sizes. In reality, *N*_*e*_ is predicted to be a lot smaller than the census sizes reported in typical E&R studies. In contrast to WFABC, CLEAR and LLS set a better standard to which one should compare new software to. Whilst CLEAR accounts for uneven coverage, LLS only considers consistency between experimental replicates. In terms of overall performance, Bait-ER and CLEAR are similar in accuracy but Bait-ER runs substantially faster. This indicates that WF and Moran models do behave similarly since MimicrEE2 simulates WF trajectories.

To investigate Bait-ER’s ability to detect selected sites in a real time series dataset, we analysed the *D. simulans* E&R experiment by Barghi et al. (2019). Bait-ER performs well on this dataset as it is rather conservative and produces only a few very significant peaks across the genome, which suggests it has a low false positive rate. It was designed to account for strong genetic drift, hence the use of a discrete-population state space. Most of the genome produced BFs greater than 2, indicating that there is not enough resolution to narrow down candidate regions to specific genes despite those very significant peaks. Barghi et al. (2019) argue that there is strong genetic redundancy caused by a highly polygenic response to selection in their experiment. Despite Bait-ER modelling sweep-like scenarios rather than the evolution of a quantitative trait using an infinitesimal model, the somewhat elevated BF signal across the genome might indicate that the genetic basis of adaptation to this new temperature regime is indeed polygenic. Our results also suggest that the impact of linked selection might be non-negligible and worth investigating further.

Linkage between selected and neutral variants has long been shown to cause skewed neutral site frequency spectra (Fay and Wu, 2000). Our analysis of the Barghi et al. (2019) experiment indicates that linked selection might be the cause of a similar skew in this dataset. Of the six genomic elements in the *D. simulans* genome, five show significant SNPs all throughout the chromosome. In fact, Buffalo and Coop (2020) have analysed temporal covariances in Barghi et al.’s dataset to quantify the impact of linked selection in a model of polygenic adaptation. They found that the covariances between adjacent time points are positive but do decay towards zero as one examines more distant time intervals. This would be predicted under a scenario where directional selection is determining the fate of linked neutral loci. Over 20% of genome-wide allele frequency changes were estimated to be caused by selection, particularly linked selection. Bait-ER producing significant hits throughout the genome is consistent with this finding. Linkage disequilibrium (LD) between neutral and selected sites is likely to have a substantial impact on genome scans such as Bait-ER that assume independence between sites.

Barghi et al. (2019) claim that their experiment showed a very distinctive pattern of heterogeneity amongst replicate populations. In other words, they found that different replicates had different combinations of alleles changing in frequency together throughout the whole experiment. In an independent study, Buffalo and Coop (2020) found that there is a substantial proportion of the initial allele frequency change in response to selection that is shared between replicates in the Barghi et al. dataset, but this pattern is overturned rapidly. This heterogeneity could be the result of at least two processes. Firstly, it can be caused by the population swiftly reaching the new phenotypic optimum, thereby hitchhiker alleles spread through the population along with adaptive sites, which are quickly fixed. These linked loci eventually experience a relaxation of selection and are free to change in frequency from then on as recombination breaks down these haplotypes. Another possibility is that such heterogeneous behaviour is not a consequence of recombination but caused by a slowdown of allele frequency changes as the population reaches the phenotypic optimum. This can leave a signature of rather distinct allele frequency spectra between replicates. A heterogeneous response can alternatively be the result of sufficient standing genetic variation followed by haplotype segregation amongst replicates. If there was enough time for the beneficial mutations to spread to different genetic backgrounds before the onset of laboratory selection, the haplotypes that were present at the foundation of the population replicates could have been segregating from the start.

The consequences of replicate heterogeneity on genome scans are twofold. On the one hand, different segregating haplotypes could be selected for in different replicates. This will cause genome scans not to find any convergent genotype frequencies, suggesting the response to selection is varied across replicates. The process is difficult to characterise unless there is sufficient data on the founder haplotypes. However, numerous studies have time series data that does not include full sequences of those starting haplotypes, e.g. Barghi et al. (2019) and Burke et al. (2014). On the other hand, if there was enough diversity at the start of the experiment, it is possible that multiple interacting beneficial mutations are already present in the standing genetic variation. Interference between linked selected sites can reduce the effectiveness of selection. This will be more prevalent if there are strongly selected sites in the vicinity. Our results indicate that that might be the case in the sweep simulated by Vlachos et al. (2019), where the authors simulate a little over 10% of the *D. melanogaster* total genome length. Each simulated segment had 30 selected targets. For moderate to strong selection, that might be enough for linkage to hinder rapid adaptation and produce signatures that are not readily captured in genome scans.

Bait-ER estimates and tests for selection. However, *σ* estimates are not to be taken literally as linked selection might be inflating individual selection coefficients. Such is the case that nearby sites are not independent from one another that whole haplotypes might be rising to fixation at once. Despite evidence that maximum likelihood estimates of selection coefficients are not affected by demography in populations as small as 500 individuals (Jewett et al., 2016), *N*_*e*_ in evolution experiments is typically even lower. Drift thus exacerbates the effect of linked selection. In addition, and in spite of researchers’ best efforts, it is common that *N*_*e*_ in laboratory experiments is lower than the census population size. For example, Barghi et al. (2019) have reared flies in populations of roughly 1000 individuals, but they have estimated *N*_*e*_ to be around 300. Therefore, in a short timescale such as that of an evolution study, recombination is unlikely to have had the chance to have broken up haplotypes present in the standing genetic variation. Collectively, our results suggest that drift should not be neglected as it might inflate selection coefficient estimates since it exacerbates the extent of linked selection. Its impact can be substantial especially for populations with low polymorphism levels.

Regardless of demographic factors, adaptation of complex traits in and of itself is a challenging process to characterise. This is because trait variation is influenced by numerous genes and gene networks. There is now some evidence in the literature suggesting that polygenic adaptation is key in a handful of laboratory evolution studies. The genomic signature left by such a complex process is still hard to describe in its entirety even in a replicated experimental design. It depends on numerous factors, including the total number of causative loci, as well as on the levels of standing genetic variation within the initial population. These are not independent of each other, as the more polygenic a trait is the more likely linkage between selected sites is to generate selected haplotypes. Nevertheless, directional selection will cause some proportion of selected sites to behave as sweep-like trajectories. It is those that Bait-ER is aiming to characterise. In short-term E&R experiments, the new phenotypic optimum could be reached towards towards the end leaving sweep signatures behind.

One aspect of time series polymorphism datasets that is worth our attention is that of missing data. It is sometimes the case that there is no frequency data at consecutive time points for a given trajectory. In the future, we will extend Bait-ER to allow for missing time points. Such a feature will enable one not to discard alleles for which not all time points have been sequenced. By using a probabilistic approach to estimate missing allele frequencies, Bait-ER has inherently the potential to cope with missing data when estimating selection parameters.

Results from genome-scans in E&R studies of small populations generally tend to underperform. Since drift is pervasive and LD is extensive, genome scans might suffer from low power and high false positive rates. For that reason, we plan to extend Bait-ER to explicitly account for linkage, which decays with distance from any given locus under selection. Accounting for linkage should help disentangle the effects of local directional selection on specific variants versus polygenic selection on complex traits. Modelling the evolution of linked sites by including information on the recombination landscape will further clarify the contribution of each type of selection.

## 4 Acknowledgements

This work was supported by the Vienna Science and Technology Fund (WWTF) through project MA16-064. CK received funding from the Royal Society (RG170315) and Carnegie Trust (RIG007474). The computational results presented have been partly achieved using the St Andrews Bioinformatics Unit (StABU), which is funded by a Wellcome Trust ISSF award (grant 105621/Z/14/Z). We are grateful to Peter Thorpe for his help with using the StABU cluster. We thank Neda Barghi and Mike Ritchie for helpful discussions, Abigail Laver and students of the BL4273 Computational Genomics module for suggestions on an early version of Bait-ER. The genome-wide scan for selected loci in the Barghi et al. (2019) dataset was conducted using the Vienna Scientific Cluster (VSC).

## Data accessibility

No new data were generated or analysed in support of this research. Bait-ER has been released as an open-source program that can be downloaded from https://github.com/mrborges23/Bait-ER, where small test datasets and a tutorial can also be found.

## Author Contributions

CB, RB and CK conceived the idea. RB developed the inferential framework and wrote the software. CB and RB tested the method on simulated data. CB performed the analyses on the Vlachos et al. (2019) and the Barghi et al. (2019) datasets. CB, CK and RB contributed to writing the manuscript.

