## Supplementary information for "Bait-ER: a Bayesian method to detect targets of selection in Evolve-and-Resequence experiments"

#### Supplementary Text

##### Impact of E&R experimental design on detecting targets of selection - sampling schemes, replication and coverage

We tested six different time schemes that allow for varying number, span, and distribution of sampled time points (schematically represented in **fig. 3A**). Surprisingly, we observed that there is no substantial increase in accuracy when more time points are sampled. Experiments with two time points estimate quite accurately $\sigma$ for most of the simulated scenarios (**fig. 3A** and **fig. S2**). Worth of note are the two time span regimes we have investigated: a span of $0.2N_e$, i.e., a common E&R study length (as seen in Barghi et al. (2019), Burke et al. (2014), and Papkou et al. (2019)), and $0.4N_e$ generations. Similarly to the number of sampled time points, we do not observe a substantial increase in the accuracy of $\sigma$ by increasing the length of the experiment.

Most E&R studies have uniform sampling schemes (Burke et al., 2014; Barghi et al., 2019). Nevertheless, we set out to compare a range of uniform and biased time schemes. The results show that there are no substantial improvements when using biased sampling schemes for neutral trajectories. However, in the presence of selection, the scheme that samples more often at the start of the experiment provides better estimates of $\sigma$ (time scheme number 3 for $N_e\sigma = 0.1$ to 10.0; **fig. 3A**).

We then set out to test the impact of the number of experimental replicate populations. We observed that the accuracy of the estimated selection coefficients increases with the number of replicated populations. The most inaccurate estimates were observed in scenarios with two replicates, both with neutral and weak selection (two replicates, $N_e\sigma = 0.0$ and 0.1; **fig. 3B**). This suggests that a higher number of replicates may help decrease the false positive rate of E&R experiments. Our results are consistent with previous studies showing that replicated populations improve the detection of selective targets (Kofler and Schlötterer, 2014) and the accuracy of estimated selection coefficients (Taus et al., 2017).

Additionally, we considered three levels of coverage to represent experiments with low to high sequencing depth. We observed that low coverage can decrease the accuracy of $\sigma$ in the case of small populations and low starting frequency trajectories (first row of the heatmap in **fig. 3C**). In contrast, the accuracy does not differ substantially in the case of average and high sequencing depths, which suggests that a coverage around 60x provides enough resolution for detecting targets of selection in E&R experiments. This is consistent with results obtained by Kofler and Schlötterer (2014), who showed that a coverage of approximately 50x is sufficient to identify strongly selected loci.

#### Real versus simulated data - comparing Barghi et al. (2019) and Vlachos et al. (2019)

To assess whether the simulation study by Vlachos et al. (2019) mimicked the experimental data gathered by Barghi et al. (2019), we examined allele frequency changes, nucleotide diversity and variance in allele frequencies amongst replicates independently for the two datasets. Overall, Barghi et al. (2019) show higher nucleotide diversity (**figs. S16** and **S17**) as well as greater allele frequency changes (**figs. S20** and **S21**). Smaller allele frequency changes are consistent with a less polymorphic population. In addition, we investigated patterns of nucleotide diversity along the chromosomes in Barghi et al. (**fig. S18**) and the genomic element simulated by Vlachos et al. (**fig. S19**). We found that both show similar regions of very low diversity at the start and/or end of the chromosome corresponding to the telomeres. In the case of Vlachos et al., the location of actual selected sites does not match dips in diversity around them, suggesting the simulation does not capture any low diversity regions that would be the result of selective sweeps. This might indicate that there is too much interference between selected sites to allow for selective sweeps or that the low polymorphism is hindering adaptation.

For both studies, we investigated the relationship between Bait-ER's scaled selection coefficient, $N_e\sigma$, and any allele frequency changes. The two show a different pattern with Barghi et al. chromosomes producing a tail of much higher $N_e\sigma$ values for trajectories that have relatively low allele frequency changes (**fig. S22**). In Vlachos et al., selected loci in yellow have allele frequency changes that range from very low to some of the highest amongst replicate populations (**fig. S23**) and we do no observe a tail of high scoring trajectories. Here, there is no apparent linear relationship regardless of replicate experiment (mean $r^2$ = -0.05, min = -0.11, max = 0.24). Moreover, we analysed how differences in coverage amongst loci would affect Bait-ER's accuracy at identifying targets of selection. This would only be an issue in Barghi et al. as all sites in the Vlachos et al. dataset have identical sequencing depth (1000x). For that reason, we visually inspected the relationship between logBFs and coverage (**fig. S24**). We found no semblance of any linear or quadratic relationship in any of the five main chromosomes, suggesting that Bait-ER is not biased towards higher coverage sites. Finally, we compared the shape of allele frequency spectra at the start (generation 0) and at the end of the experiment (generation 60) for both studies (**figs. S25** and **S26**). Barghi et al. spectra are markedly U-shaped with a non-negligible amount of intermediate frequency alleles. In contrast, Vlachos et al. simulations show distribution that is much more biased towards low frequency alleles which is compatible with the previous finding of low nucleotide diversity in this dataset.

### Supplementary tables and figures

| Conditions | Relative CPU time |
|---|---|
| Base experiment | 1.00 |
| 2 replicates | 0.97 |
| 10 replicates | 1.15 |
| 100 individuals | 0.05 |
| 1000 individuals | 31.97 |
| 20X coverage | 1.00 |
| 100X coverage | 1.00 |
| 2 time points | 0.92 |
| 11 time points | 1.16 |

Table S1: **Relative CPU time for Bait-ER under several experimental/population conditions.**[a]

[a]The relative CPU time was calculated considering a base E&R experiment with 5 replicates, 5 uniformly distributed time points, 300 individuals and an average coverage of 60X. Unless otherwise specified, the remaining simulation parameters were the same as those under base experiment conditions.

| Parameter | | Simulated values | Notes |
|---|---|---|---|
| **Population parameters** | | | |
| $N_e$ | effective population size | 100, **300** and 1000 | representing a small, a typical and a large in E&R study population |
| $p_0$ | allele's initial frequency | **0.01, 0,05, 0.1 and 0.5** | representing rare, low frequency and common alleles |
| $\sigma$ | selection coefficient | $\mathbf{0, 1/10N_e, 1/N_e}$ **and** $\mathbf{10/N_e}$ | representing regimes of neutrally evolving, drift-dominated, and selection-dominated allele trajectories |
| **Experimental parameters** | | | |
| $C$ | Coverage | 20x, **60x** and 100x | low, medium and high coverage for pool-seq data |
| $R$ | Number of replicates | 2, **5** and 10 | |
| $T$ | Number of time points | 2, **5** and 11 time points, assessed at generations<br><br>(0.0, 0.2),<br>**(0.00, 0.05, 0.10, 0.15, 0.20)**,<br>(0.00, 0.04 0.08 0.12 0.20),<br>(0.00, 0.08 0.12 0.16 0.20),<br>(0.0, 0.1 0.2 0.3 0.4) and<br>(0.00, 0.02 0.04 0.06 0.08 0.10 0.12 0.14 0.16 0.18 0.20) relative to $N_e$. | represents different combinations of total number of time points, experiment lengths<br>and distribution of sampling events (uniform/non-uniform) |

Table S2: **Simulated scenarios.**[b]

[b]The simulated parameters can be divided into two categories: those which are related with the population dynamics (effective population size, selection coefficient, and allele's starting frequency) and those related to the experimental design (coverage, number of time points and number of replicates). To test the experimental conditions, we defined a base experiment with 5 replicates, 5 uniformly distributed time points (total span of $0.20Ne$ generations) and a coverage of 60x. This base experiment is highlighted in bold. The two maximum experiment lengths considered ($0.2N_e$ and $0.4N_e$) were chosen based on typical E&R experimental designs.

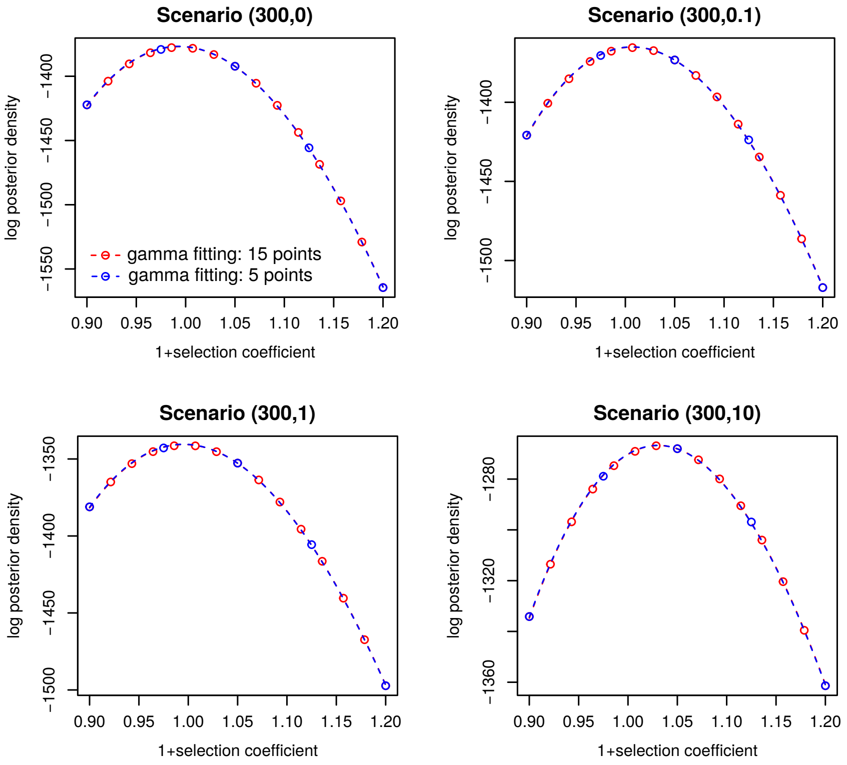


Figure S1: **Gamma fit to the log posterior distribution of $\sigma$.** The fitting of the gamma density to the posterior distribution of $1+\sigma$ was evaluated for both neutral and selected trajectories. Each plot represents the different scenarios that were simulated by varying the population size, selection coefficient (indicated within brackets ($N_e$, $N_e\sigma$)). The points represent either 15 or 5 assessments of the posterior distribution (red and blue points, respectively) obtained whilst computing the log-likelihood. The dashed lines represent the gamma fitting to the log posterior obtained by using either 15 or 5 points. Both fitting methods match quite well, suggesting Bait-ER's current approach using 5 points is a good choice.

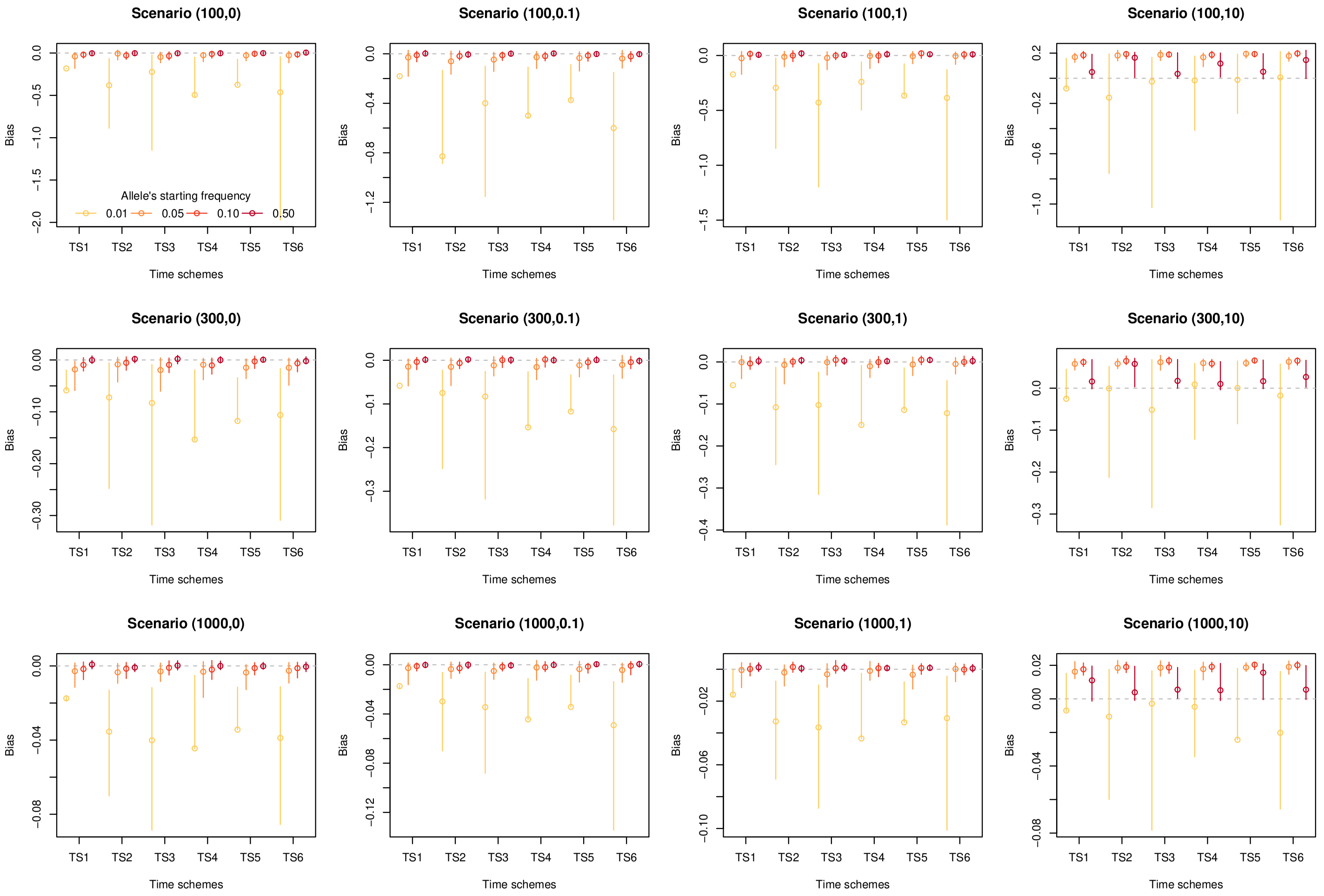


Figure S2: **Impact of the number and distribution of sampled time points, as well as experimental span on the estimated selection coefficients.** The plots illustrate the bias of the estimated selection coefficients for different time sampling schemes of an E&R experiment. This bias was calculated by computing the difference between the Bait-ER estimated and the true $\sigma$: $\hat{\sigma} - \sigma$. The vertical lines and points indicate the interquartile range and median bias. Each plot represents a scenario that was simulated by varying the population size, the true selection coefficient (indicated within brackets ($N_e$, $N_e\sigma$)) and starting allele frequency (indicated by the yellow-to-red colour gradient). Here is a brief explanation of how the different time schemes might vary from one another: most time schemes have five sampling times, apart from TS1 and TS6, which had two and eleven sampling points, respectively; the majority of time schemes have a total span of $N_e/5$ generations, except for TS5 which had double the span ($2N_e/5$); lastly, uniform sampling was used across most scenarios, but for TS3, which is more heavily sampled during the first half of the experiment, and TS4, during the second half.

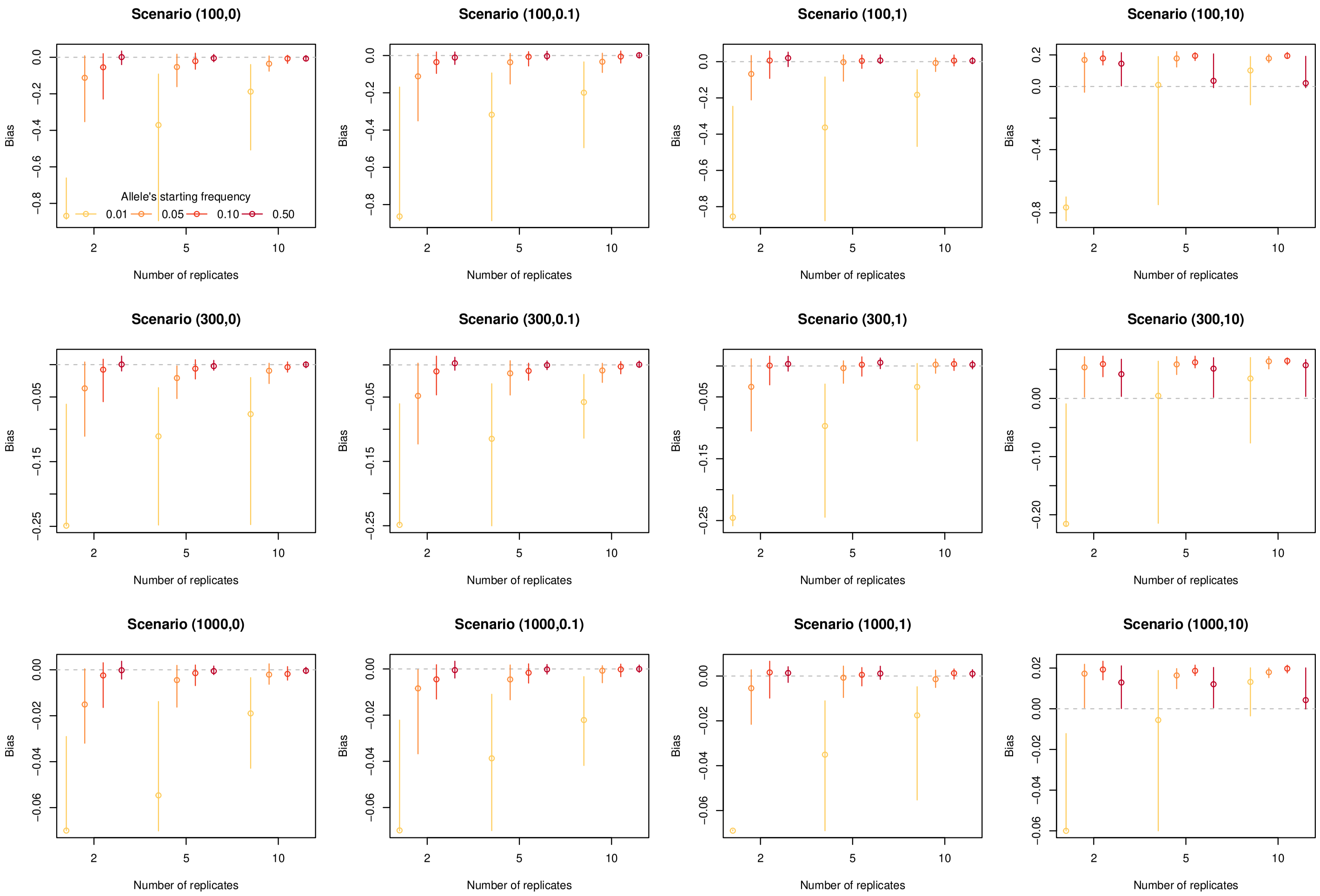


Figure S3: **Impact of the number of replicates on the estimation of selection coefficients.** The plots show the bias of the estimated selection coefficients for several scenarios with different number of replicate experimental populations. This bias was calculated by computing the difference between the Bait-ER estimated and the true $\sigma$: $\hat{\sigma} - \sigma$. Vertical lines and points indicate the interquartile range and median bias. Each plot represents the different scenarios that were simulated by varying the population size, the true selection coefficient (indicated within brackets ($N_e$, $N_e\sigma$)) and starting allele frequency (indicated by the yellow-red gradient of colors).

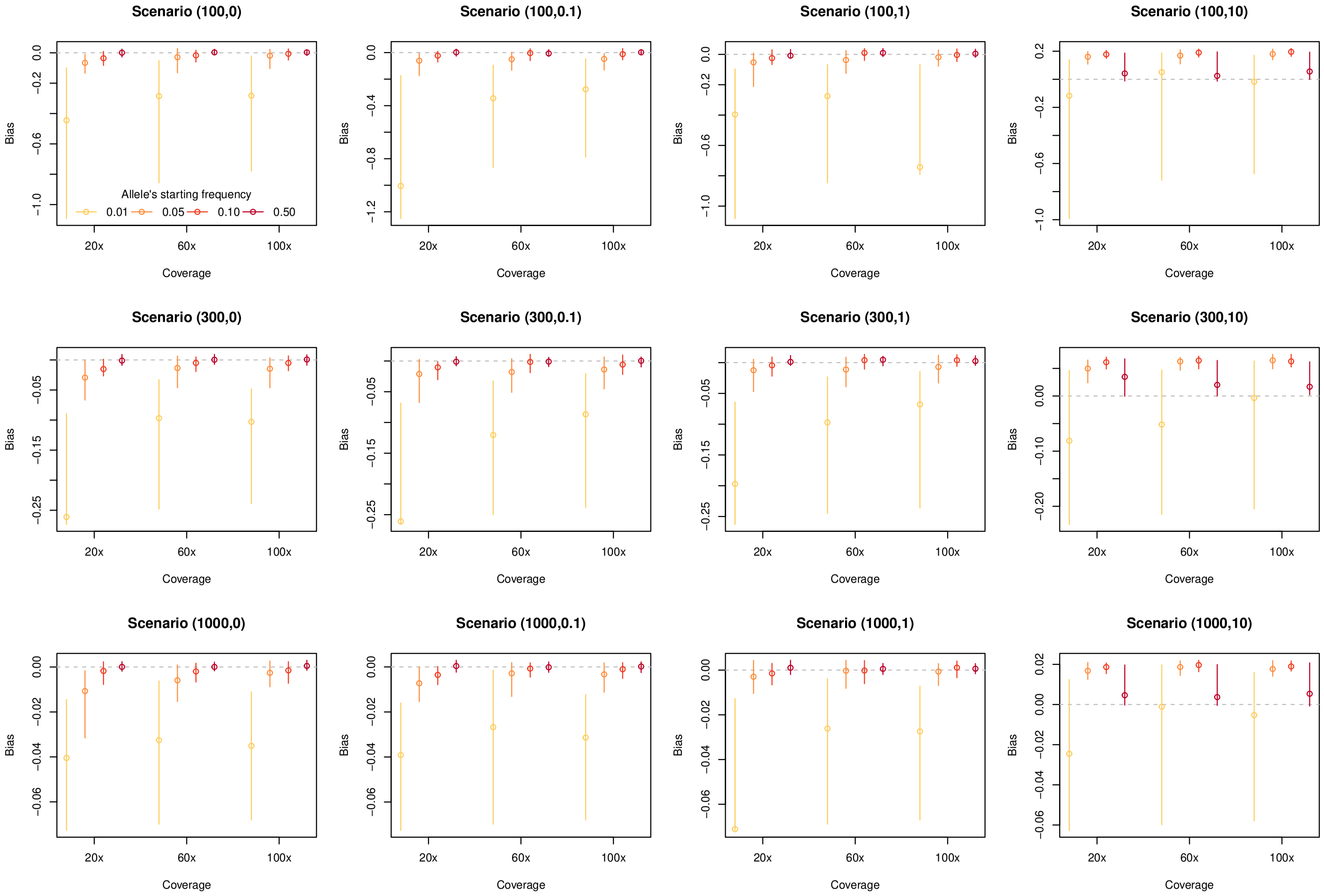


Figure S4: **Impact of coverage on the estimation of selection coefficients** The plots show the bias of the estimated selection coefficients obtained for different sequencing depths of an E&R experiment. This bias was calculated by taking the difference between the Bait-ER estimated and the true $\sigma$: $\hat{\sigma} - \sigma$. The vertical lines and points indicate the interquartile range and median bias. Each plot represents the different scenarios that were simulated by varying the population size, selection coefficient (indicated within the brackets ($N_e$, $N_e\sigma$)) and starting allele frequency (indicated by the yellow-to-red colour gradient).

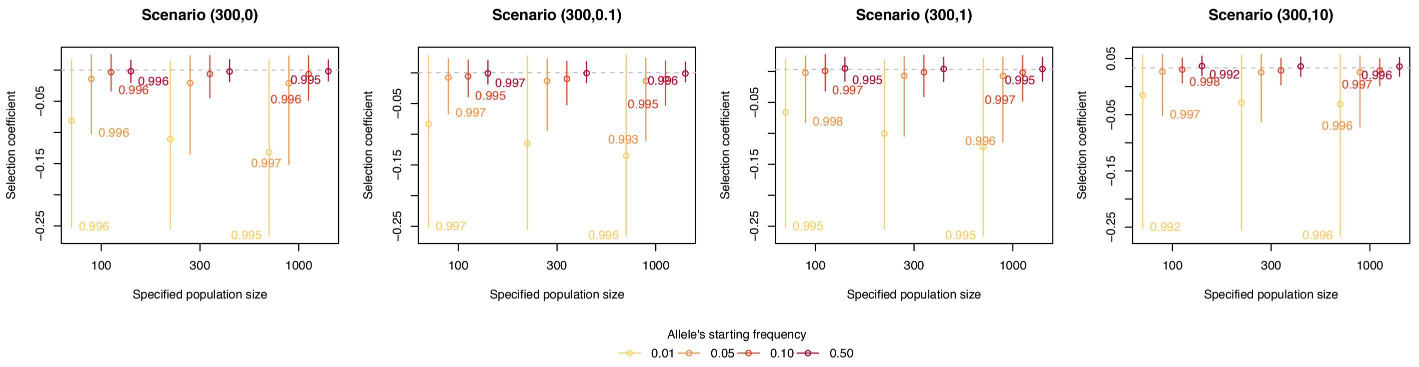


Figure S5: **Impact of the user-specified population size on the estimation of selection coefficients.** The plots show the distribution of the estimated selection coefficients where the population size is misspecified. Vertical lines and points indicate the interquartile range and median selection coefficient. Each plot represents a specific scenario that was simulated by varying the population size, the true selection coefficient (indicated within brackets ($N_e, N_e\sigma$)) and starting allele frequency (indicated by the yellow-to-red colour gradient). The numbers next to each bar correspond to the Spearman's correlation coefficient, which correlate the BFs of the 100 replicated trajectories between the cases where we have either under- and overspecified the population size ($N_e = 100$ or 1000, respectively) and the case where we use the true population size ($N_e = 300$).

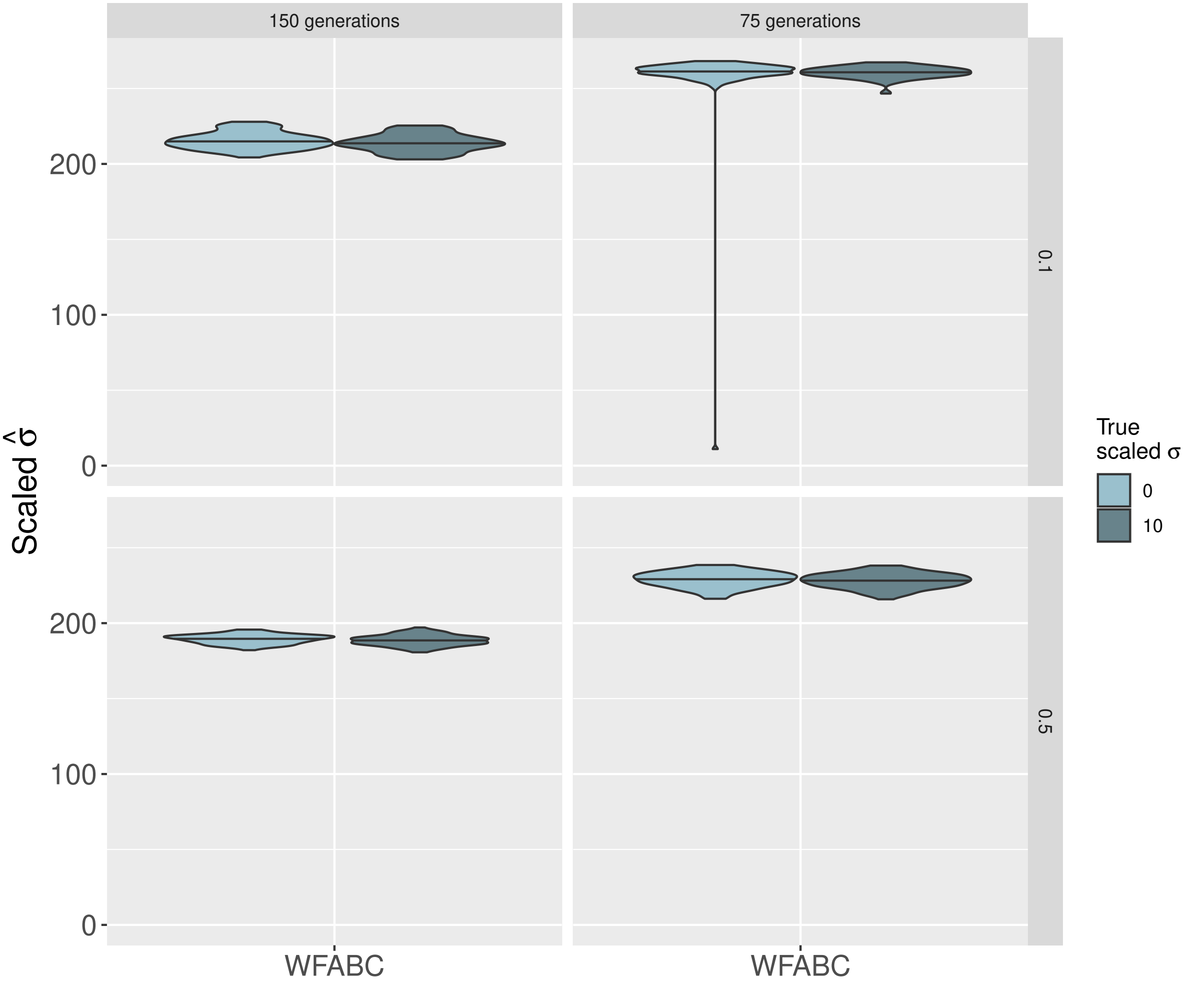


Figure S6: **Comparison of estimates of $\sigma$ produced by WFABC**. These plots include estimates for those Moran trajectories simulated with starting frequencies of 10% and 50% (top and bottom row, respectively). Only neutrally evolving ($N_e\sigma = 0$) and strongly selected alleles were considered here ($N_e\sigma = 10$). The left and right hand side panels correspond to two different experiment lengths: 150 and 75 generations, respectively.

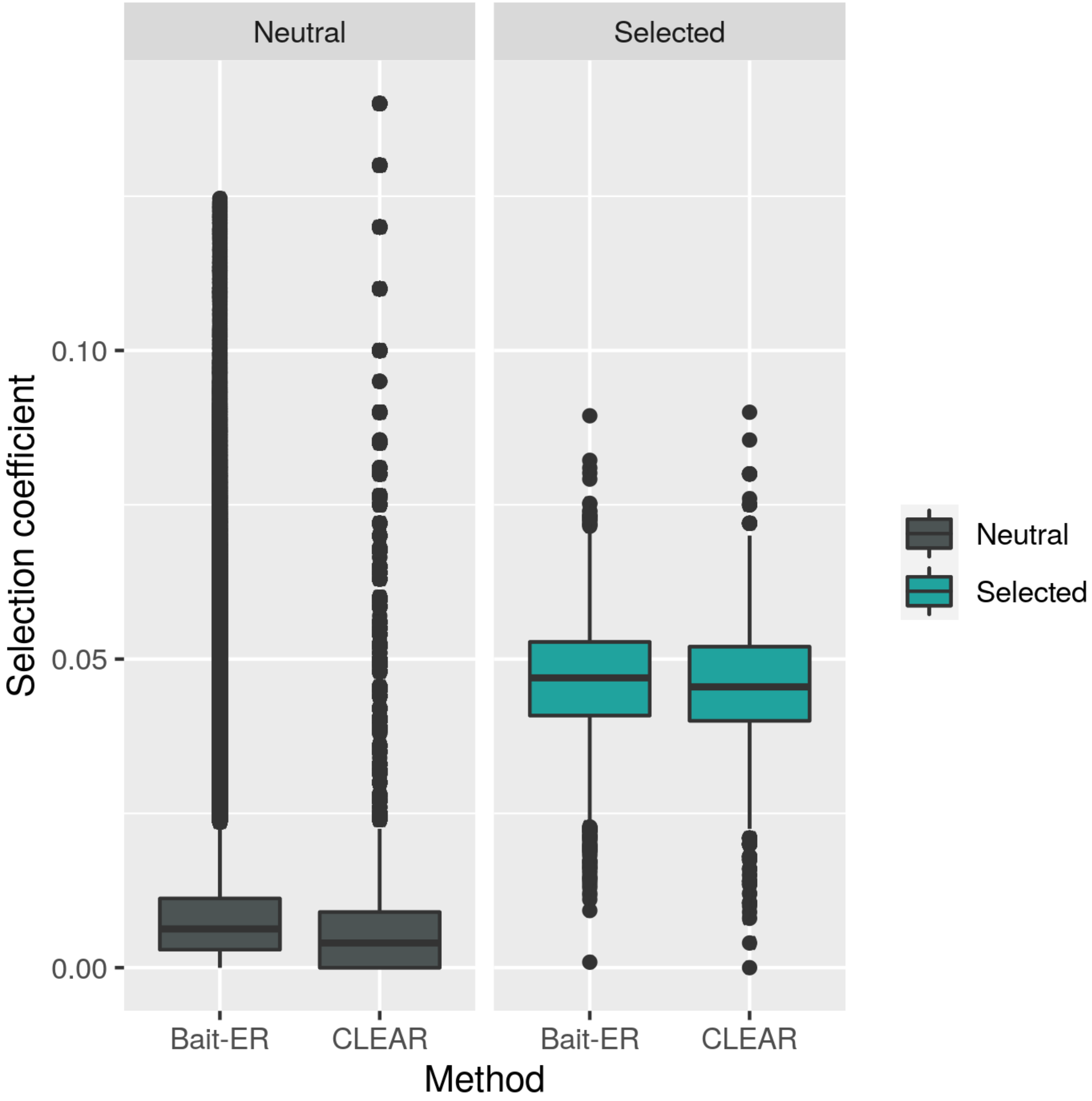


Figure S7: **Distribution of $\hat{\sigma}$ for Bait-ER and CLEAR.** These boxplots show the comparison between estimates of selection coefficients for both selected (right panel) and neutral (left panel) sites from the Vlachos et al. (2019) selective sweep scenario. Data filtered out by Bait-ER was excluded from the analysis.

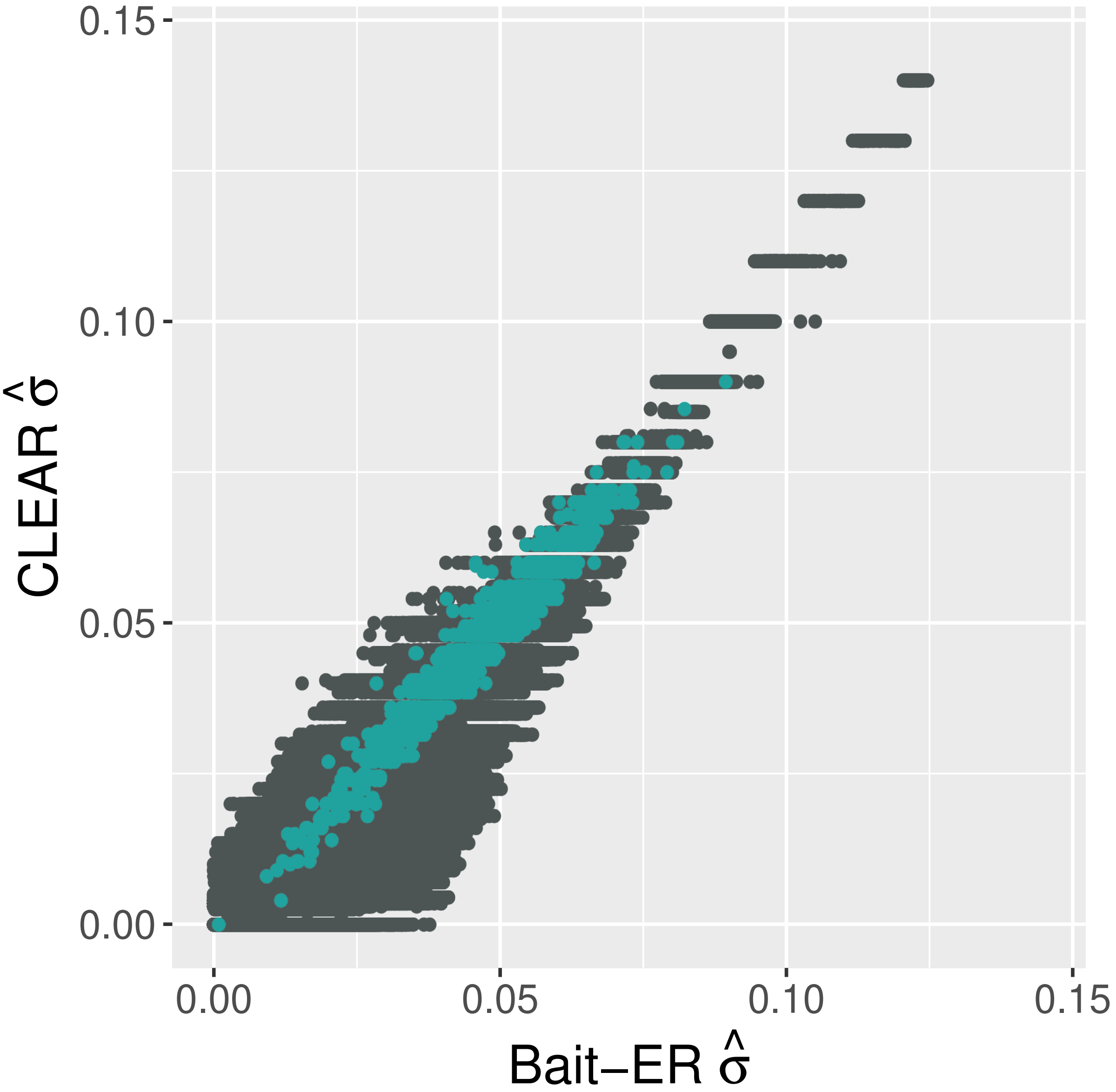


Figure S8: **Comparison of Bait-ER vs CLEAR $\hat{\sigma}$ in the Vlachos et al. sweep scenario.** This scatterplot is the comparison of selection coefficient estimates for both methods, where blue and grey points are selected and neutral sites, respectively. Data filtered out by Bait-ER was excluded from the analysis.

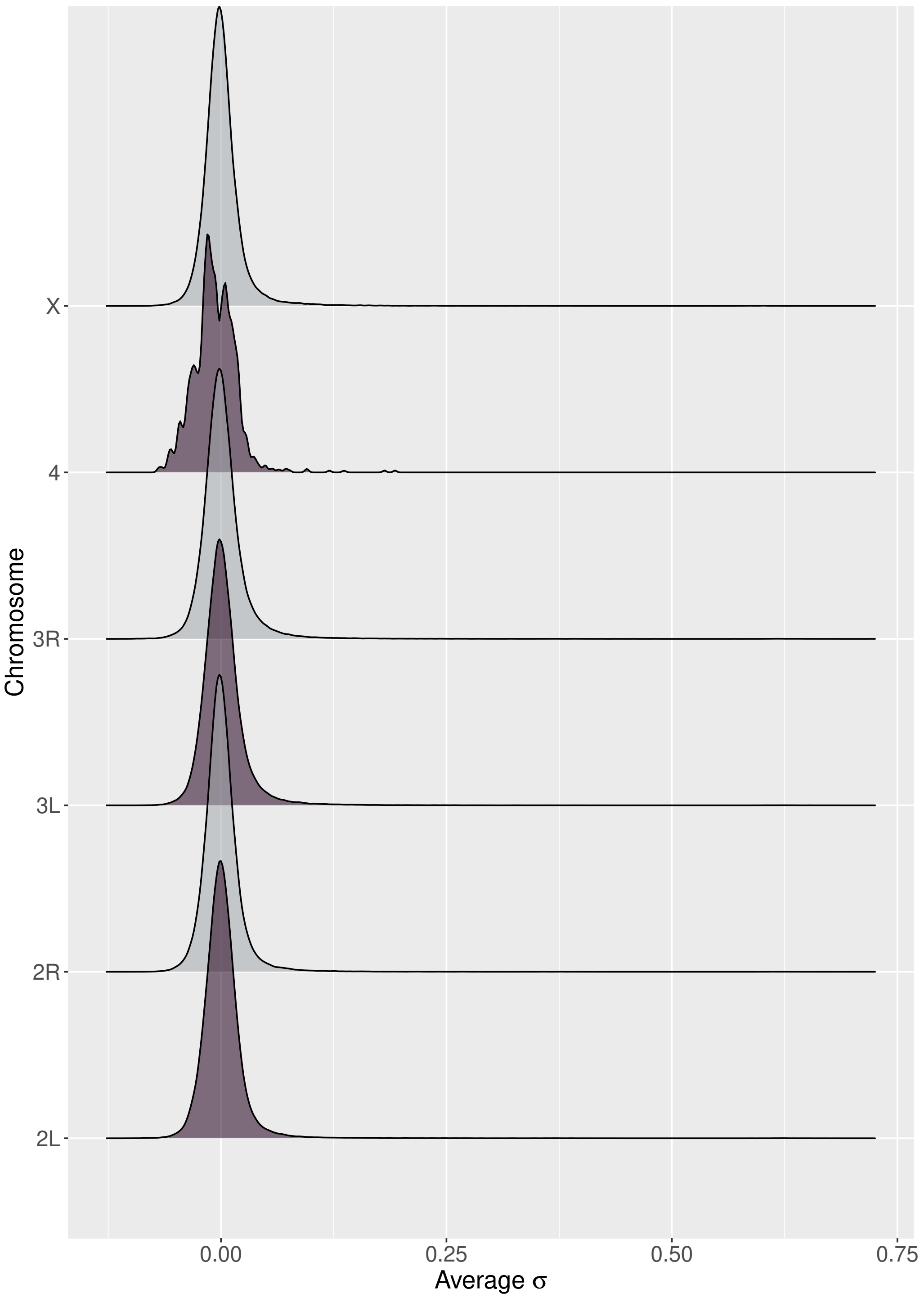


Figure S9: **Distribution of the selection coefficients estimated by Bait-ER for each of the chromosomes in the Barghi et al. (2019) dataset.** From bottom to top row, the figure shows the distribution of $\hat{\sigma}$ on chromosomes 2L, 2R, 3L, 3R, 4 and X. All distributions seem to be centered around 0, which is unsurprising given that we expect selection not to be widespread across the genome but rather restricted to some genomic windows rather.

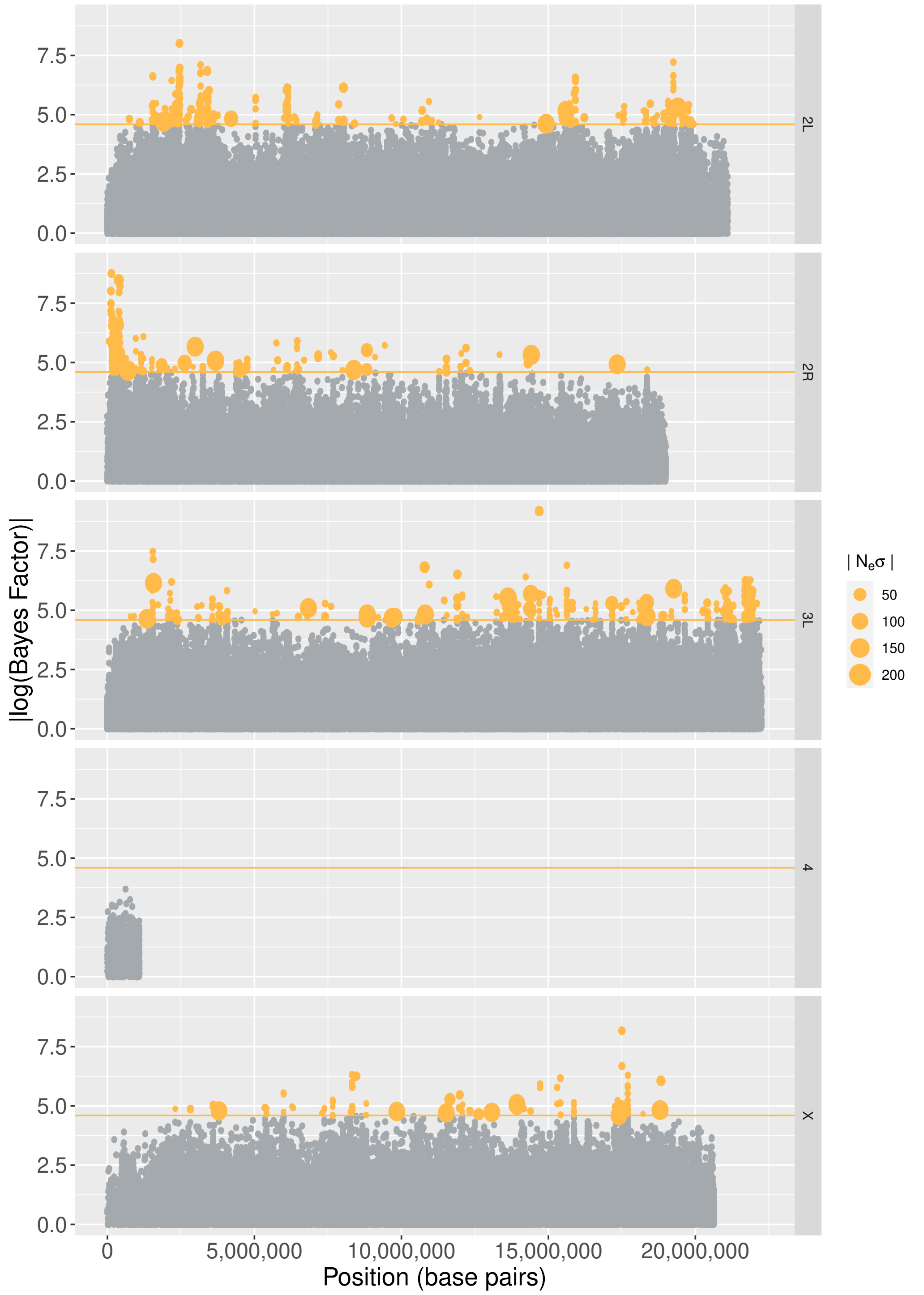


Figure S10: **Bayes Factors on chromosomes 3L, 2, 4, 5 and X.** Similarly to figure 8, these Manhattan plots show log-transformed Bayes Factors computed by Bait-ER for loci along the left arm of the 3$^{rd}$ chromosome, as well as chromosomes 2, 4, 5 and X in the Barghi et al. (2019) time series dataset. The orange line indicates a conservative threshold of approximately 4.6, which corresponds to log(0.99/0.01), meaning all points in orange have very strong evidence for these to be under selection. The SNPs that are significant at this level are sorted by size according to how strong Bait-ER's selection coefficients are. In other words, points are sized according to how strong the large selection coefficient is estimated to be.

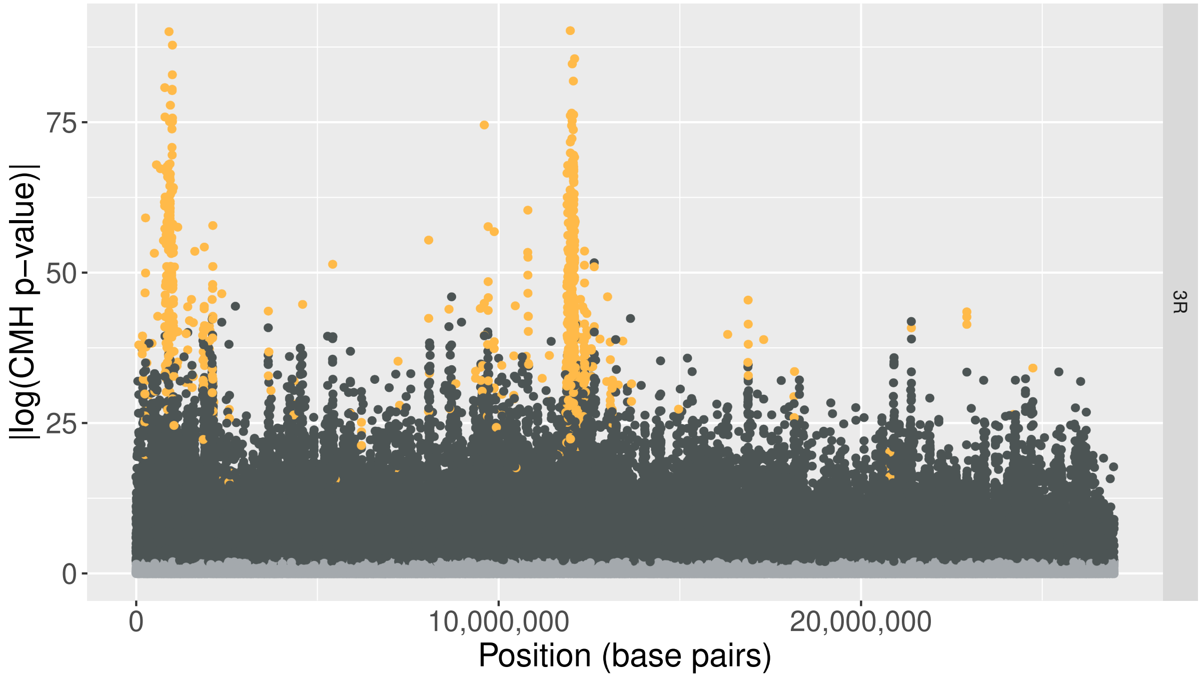


Figure S11: **CMH test p-values along chromosome 3R.** Data presented are the log-transformed p-values produced by the CMH test as per Barghi et al. for comparison with fig. 8. The most pronounced peaks seem to be common to both methods, although the CMH test produces many more significant values when compared to Bait-ER. Orange coloured points correspond to BFs which are greater than log(0.99/0.01) (approx. 4.6) and p-values less than or equal to 0.01, i.e., those that are considered significant by both tests. Blue coloured points indicated that the computed BF is greater than our threshold but not significant according to the CMH test. Additionally, dark grey points are significant according to the CMH test, but not to Bait-ER, and light grey points are inferred not significant by both tests.

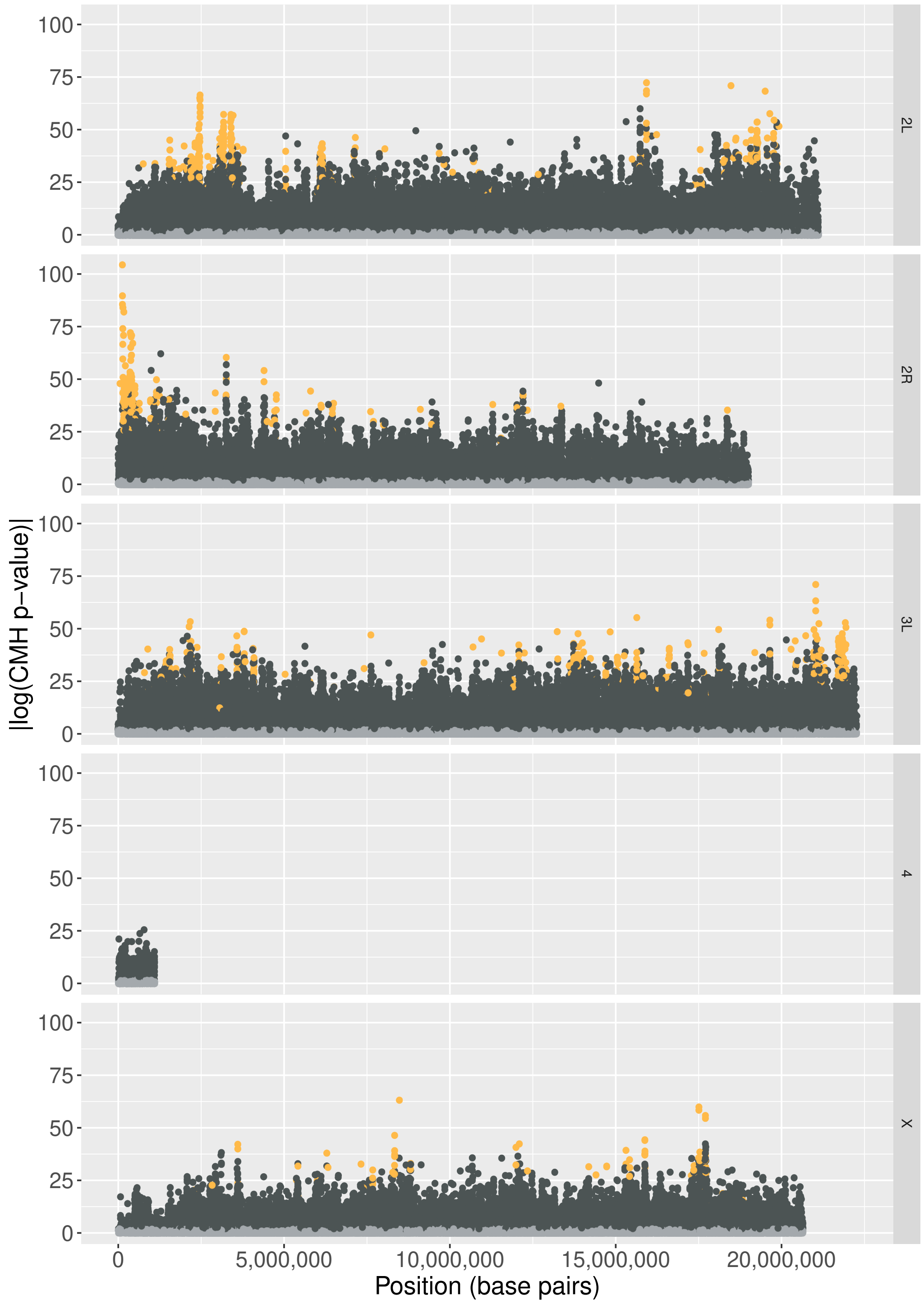


Figure S12: **CMH test p-values along chromosomes 3L, 2, 4, 5 and X.** Data presented are the log-transformed p-values produced by the CMH test as per Barghi et al. for comparison with fig. S10. Orange coloured points correspond to BFs which are greater than log(0.99/0.01) (approx. 4.6) and p-values less than or equal to 0.01, i.e., those that are considered significant by both tests. Blue coloured points indicated that the computed BF is greater than our threshold but not significant according to the CMH test. Additionally, dark grey points are significant according to the CMH test, but not to Bait-ER, and light grey points are inferred not significant by both tests.

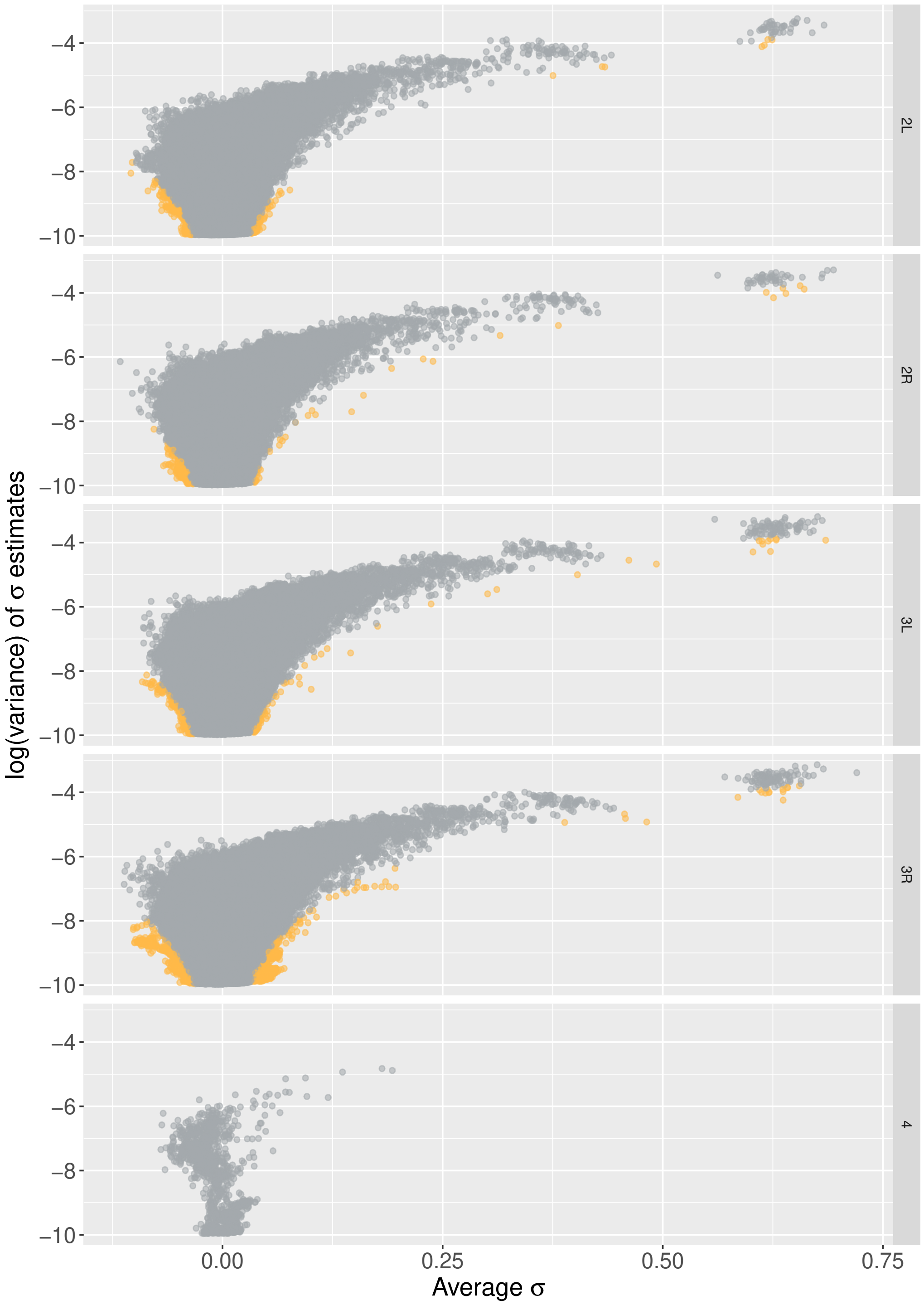


Figure S13: **Variance versus mean sigma on chromosomes 3R, 3L, 4 and 5.** Similarly to fig. 9, these graphs compare log transformed variances in $\sigma$ estimates to average $\sigma$s. The variance is calculated using the inferred rate and shape parameters for the beta distribution, and the average $\sigma$ is the mean value of the posterior distribution estimated by Bait-ER. Orange coloured points are significant at a conservative BF threshold of log(0.99/0.01), approx. 4.6. Data in Barghi et al. (2019).

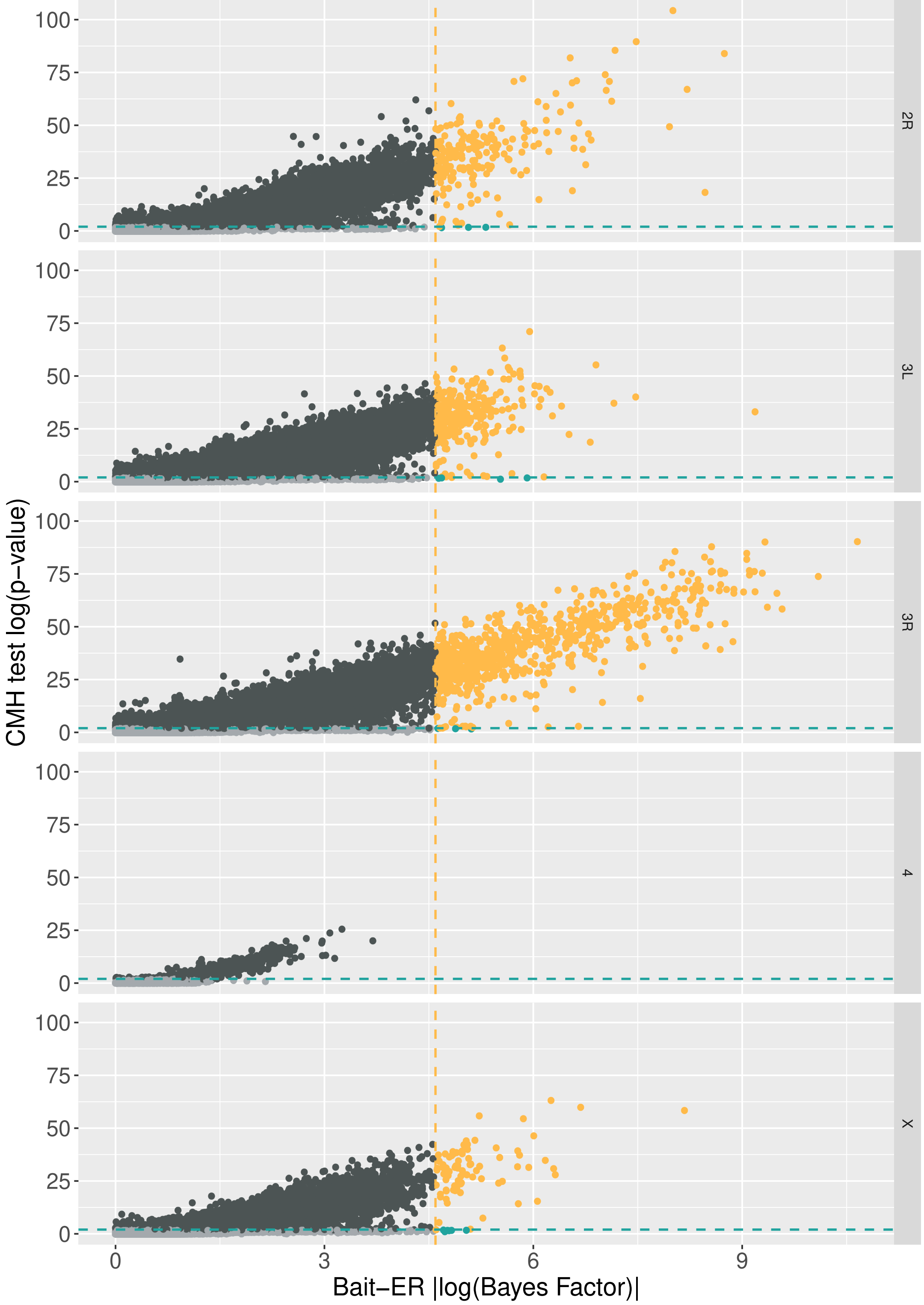


Figure S14: **Bait-ER's Bayes Factors versus CMH test's p-values on chromosomes 2L, 3R, 3L, 4 and X.** Orange coloured points correspond to BFs which are greater than log(0.99/0.01) (approx. 4.6) and p-values less than or equal to 0.01, i.e., those that are considered significant by both tests. Blue coloured points indicated that the computed BF is greater than our threshold but not significant according to the CMH test. Additionally, dark grey points are significant according to the CMH test, but not to Bait-ER, and light grey points are inferred not significant by both tests.

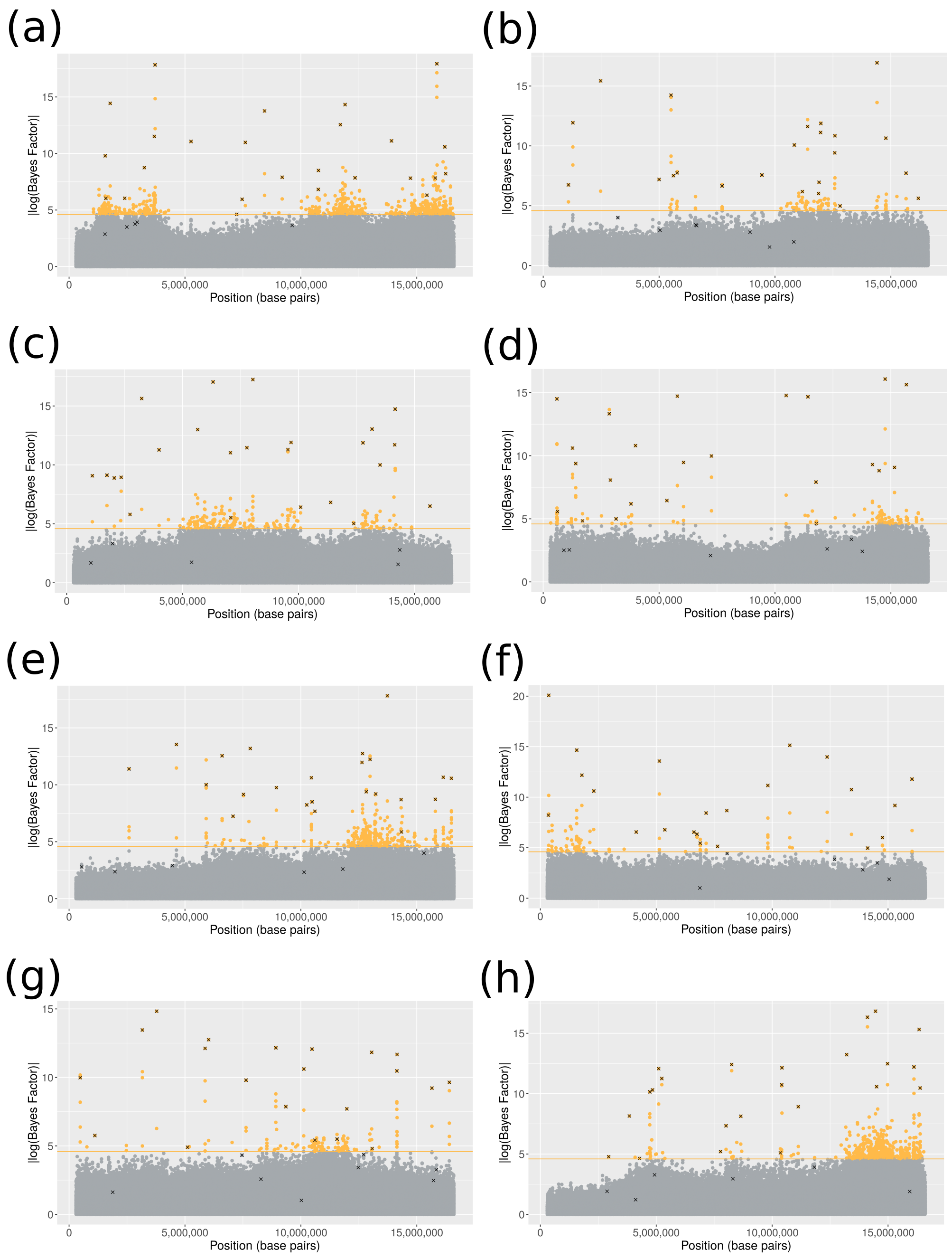


Figure S15: **Bayes Factors on 8 replicates experiments on the chromosome arm simulated in Vlachos et al. (2019).** This Manhattan plot shows log-transformed Bayes Factors computed by Bait-ER. The orange line indicates a conservative threshold of approximately 4.6, which corresponds to log(0.99/0.01). Data points which are marked by an x are those that were the true targets of selection. These were randomly chosen and simulated with a constant selection coefficient of 0.5. Plots a-h correspond to replicate experiments #4, #9, #39, #50, #51, #77, #84 and #93, respectively.

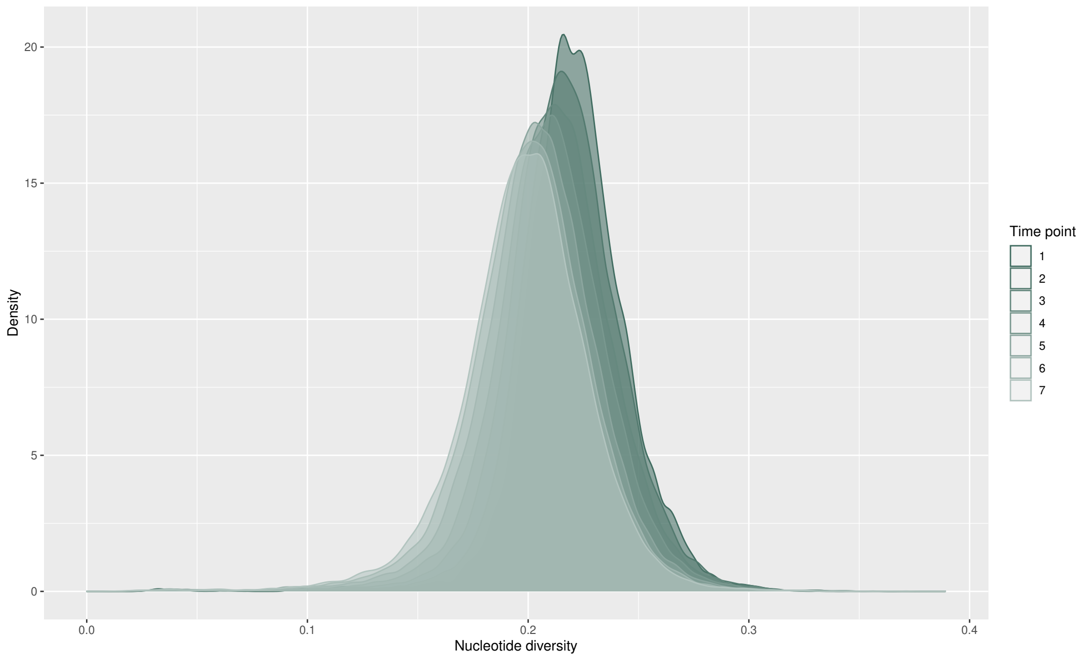


Figure S16: **Relative nucleotide diversity density per time point for all chromosomes (excluding 4) in the Barghi et al. (2019) dataset.** Diversity values presented are per site.

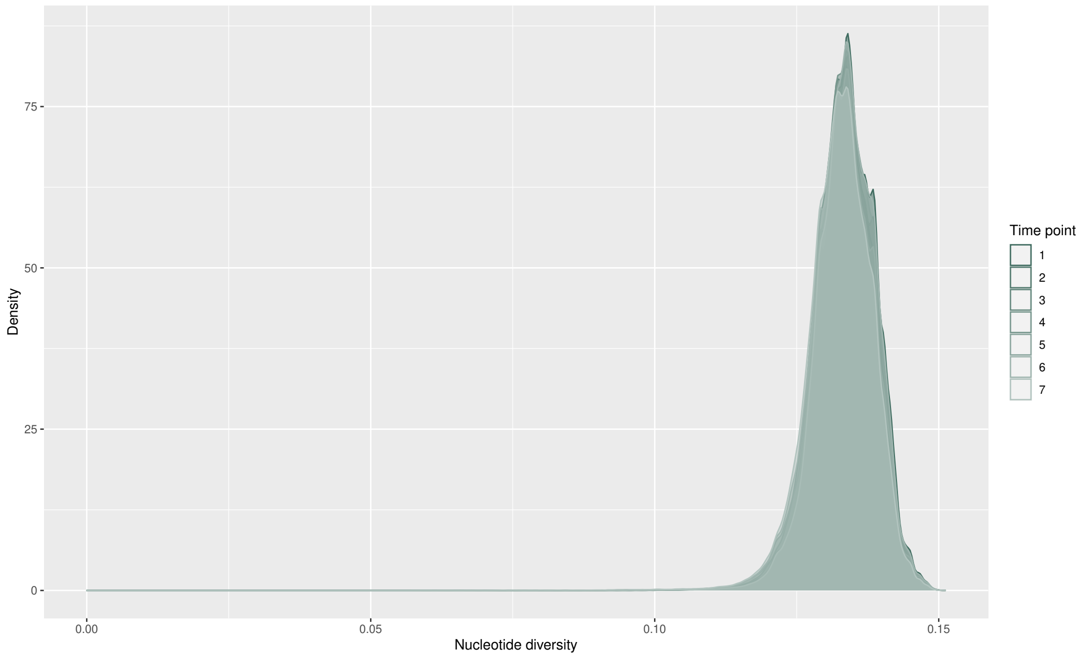


Figure S17: **Relative nucleotide diversity density per time point for all replicate experiments and populations in the Vlachos et al. (2019) dataset.** Diversity values presented are per site.

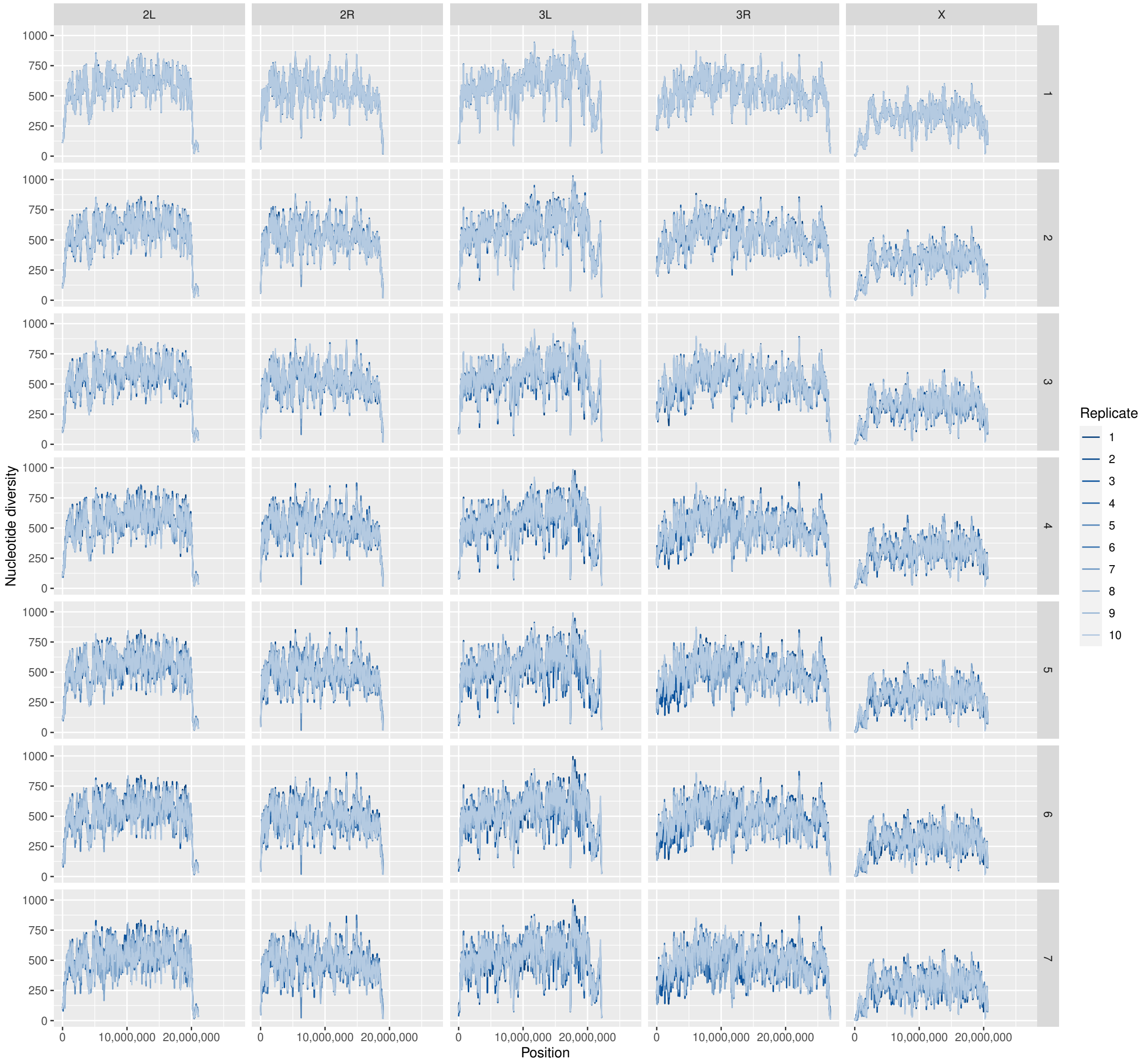


Figure S18: **Absolute nucleotide diversity along each chromosome (excluding 4) in the Barghi et al. (2019) dataset.** Diversity values presented are per window. Rows correspond to consecutive time points.

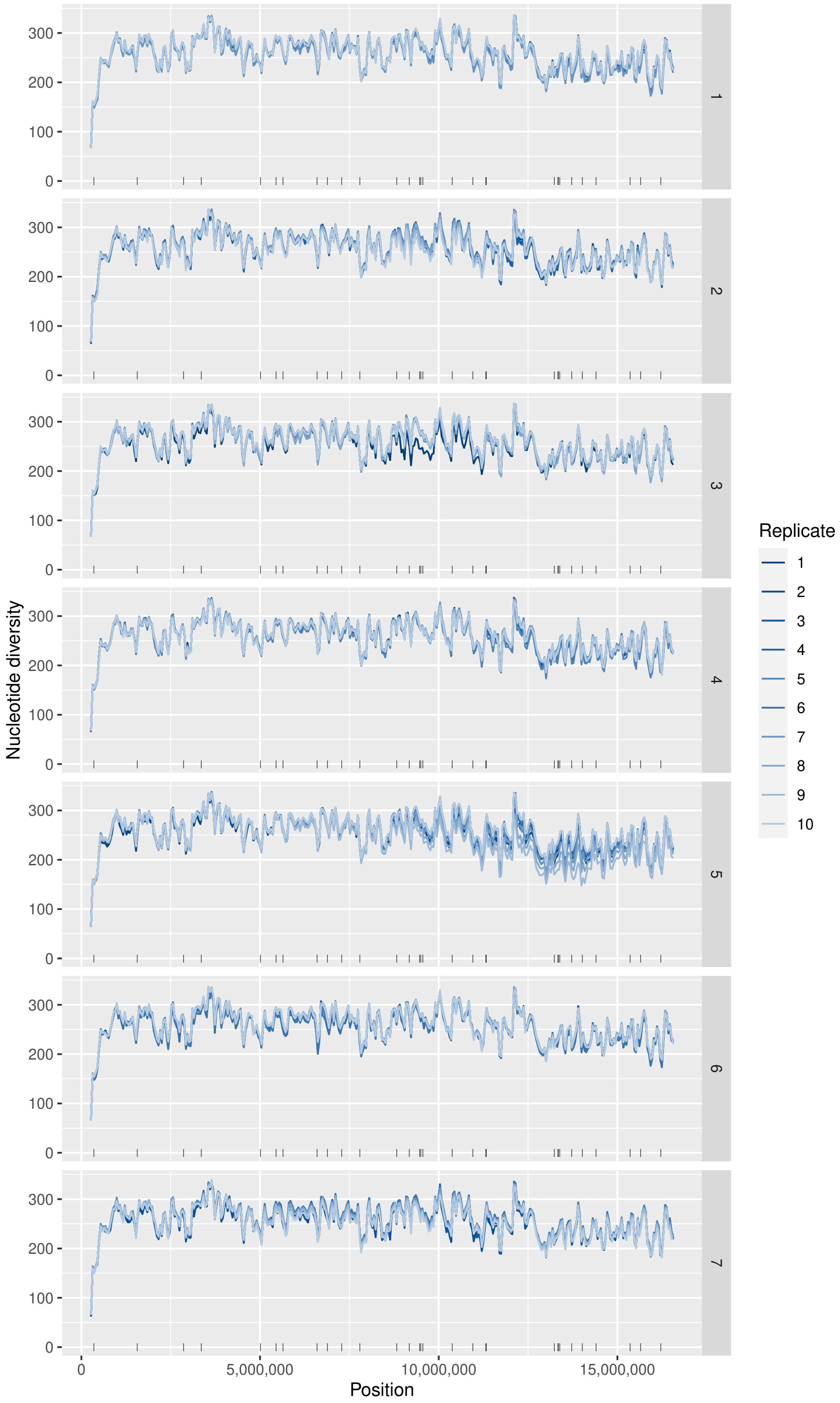


Figure S19: **Absolute nucleotide diversity along the genomic element simulated by Vlachos et al. (2019) – replicate experiment #18.** Diversity values presented are per window. Rows correspond to consecutive time points. Black solid bars at the bottom of each individual plot indicate the location of a selected site.

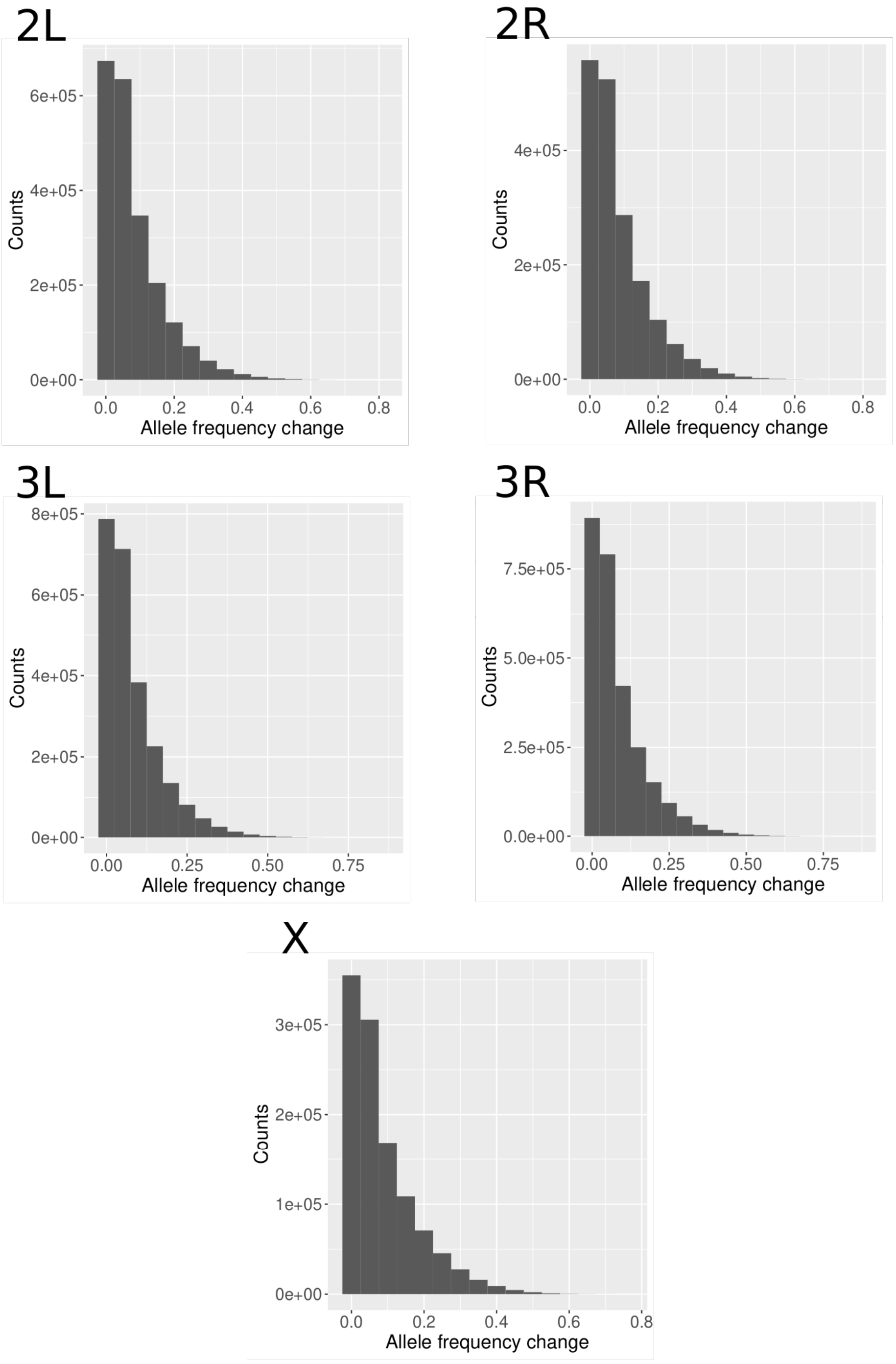


Figure S20: **Histogram of total allele frequency changes in Barghi et al. (2019) chromosomes.** Excludes data on the fourth chromosome.

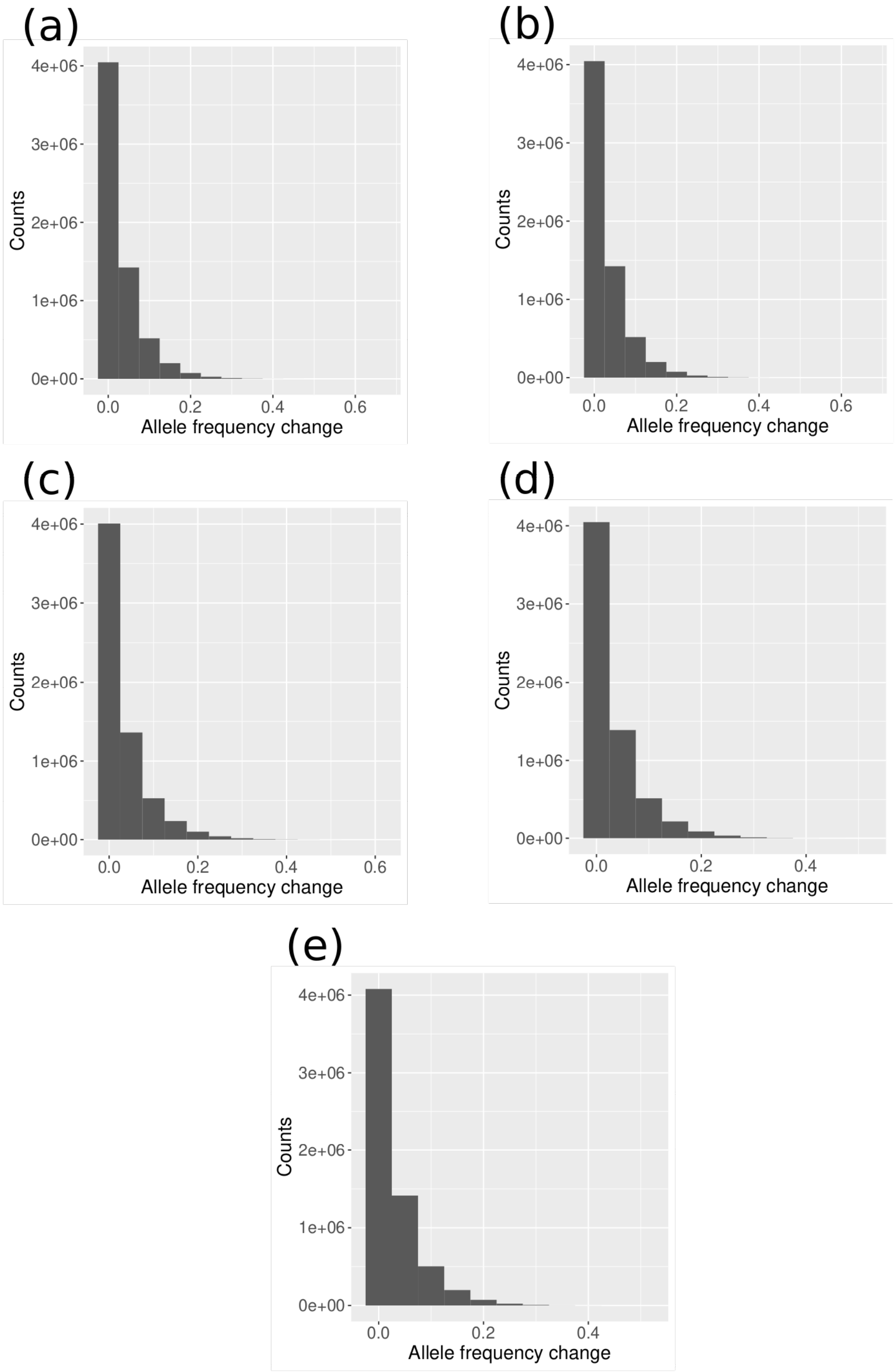


Figure S21: **Histogram of total allele frequency changes in five Vlachos et al. (2019) sweep scenario experiments.** Includes data on replicate experiments (a) #2, (b) #7, (c) #49, (d) #76 and (e) #100.

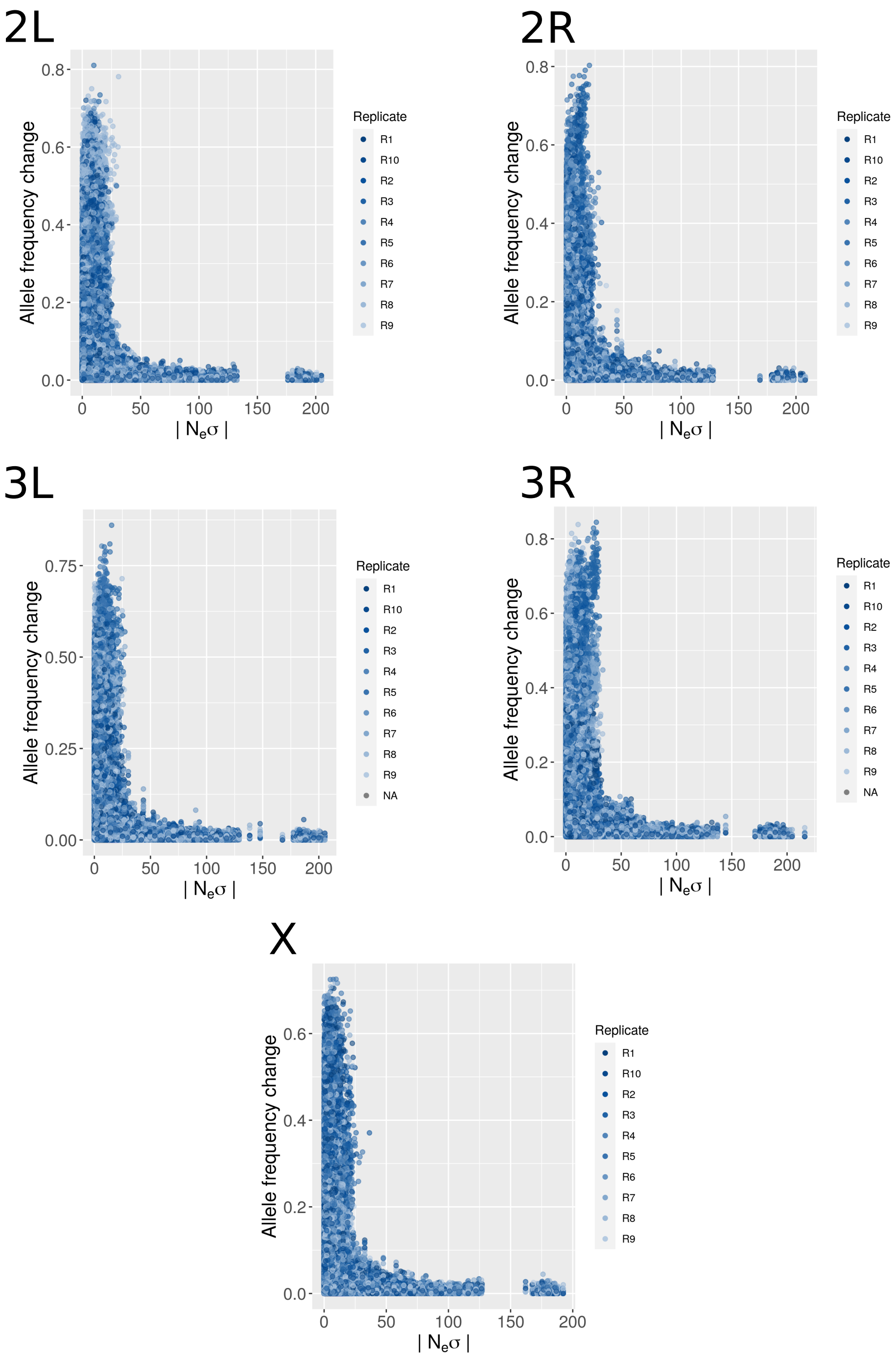


Figure S22: **Total allele frequency changes for each locus in the Barghi et al. (2019) dataset versus $N_e\sigma$.** Excludes data on the fourth chromosome.

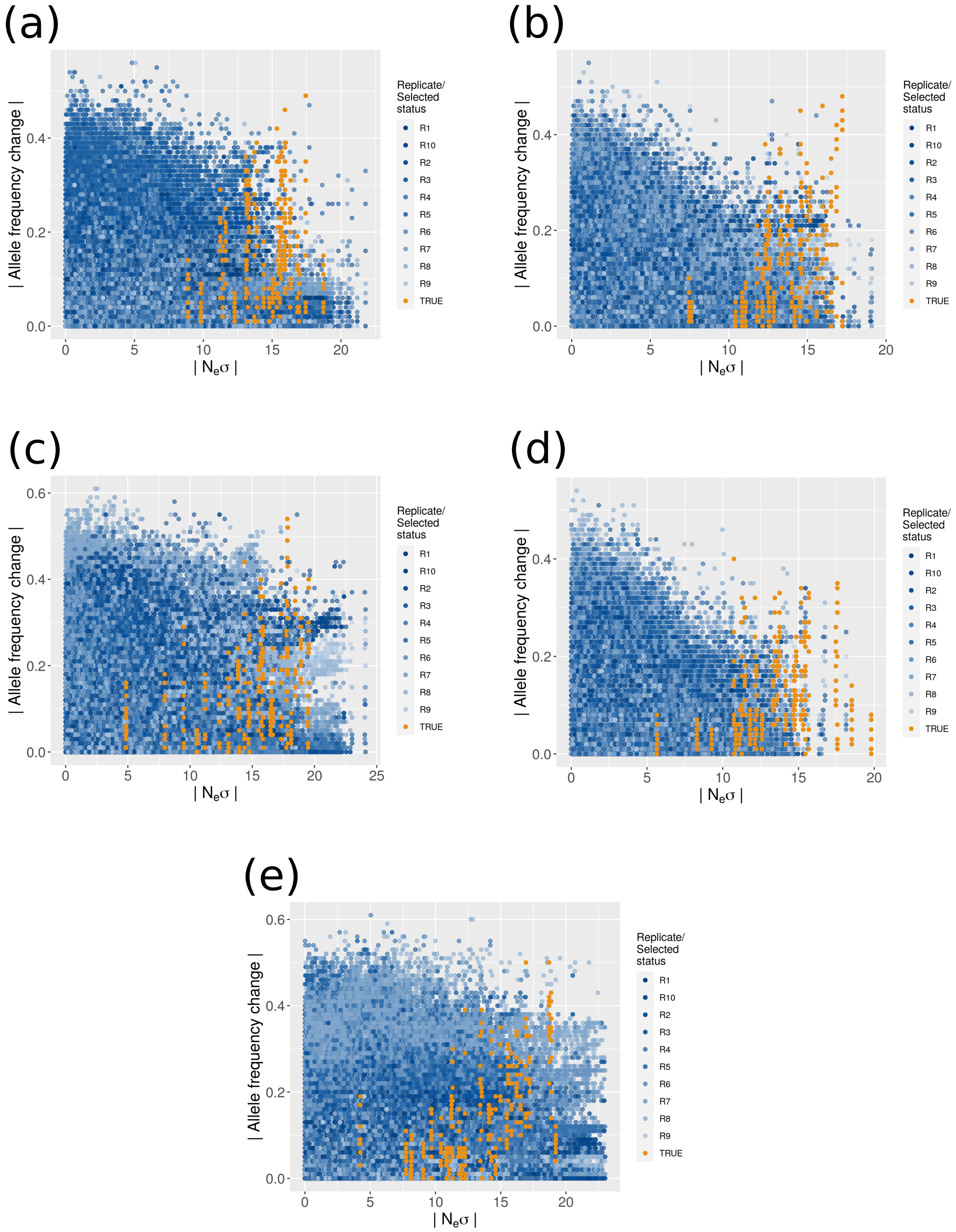


Figure S23: **Total allele frequency changes for each locus in the Vlachos et al. (2019) dataset versus $N_e\sigma$.** Includes data on replicate experiments (a) #31, (b) #31, (c) #47, (d) #54 and (e) #76. Orange points indicate true selected sites.

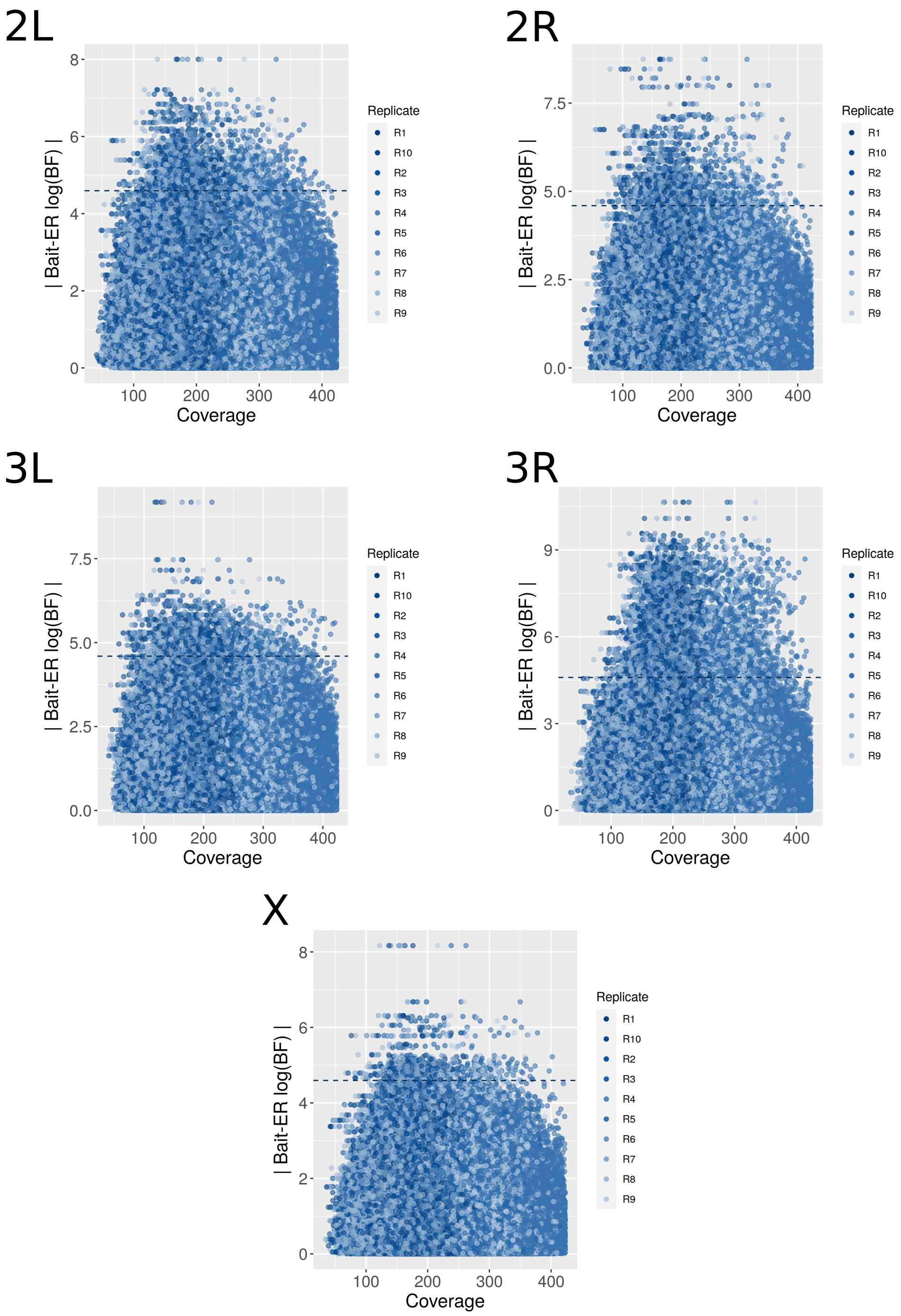


Figure S24: **Scatterplot of the relationship between coverage at generation 0, i.e. time point 1, and Bait-ER logBFs for chromosomes in the Barghi et al. (2019) dataset.** Excludes data on the fourth chromosome.

Generation 0

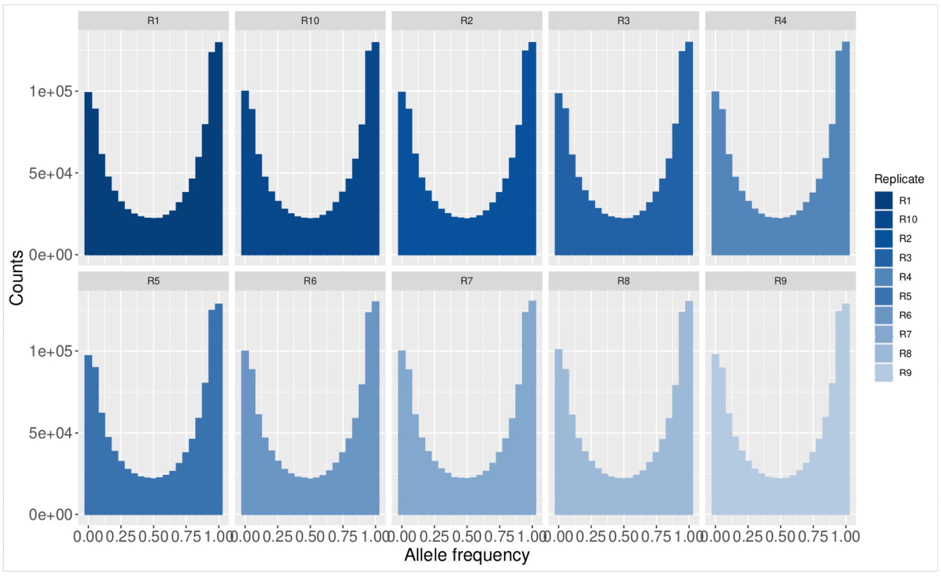


Generation 60

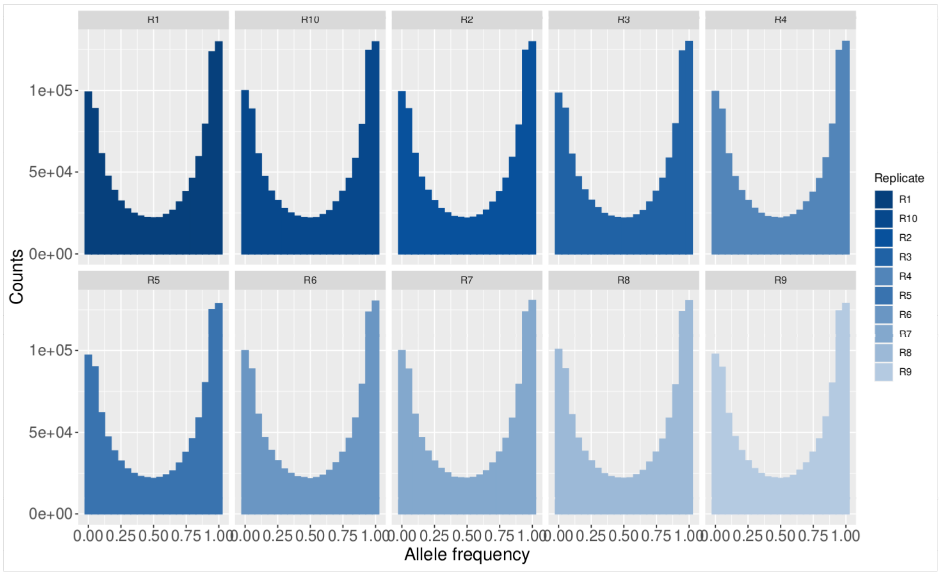


Figure S25: **Allele frequency spectra of chromosome 2L in Barghi et al. (2019) at generations 0 (top) and 60 (bottom), i.e., time points 1 and 7.** Each replicate population is represented in its own individual histogram plot.

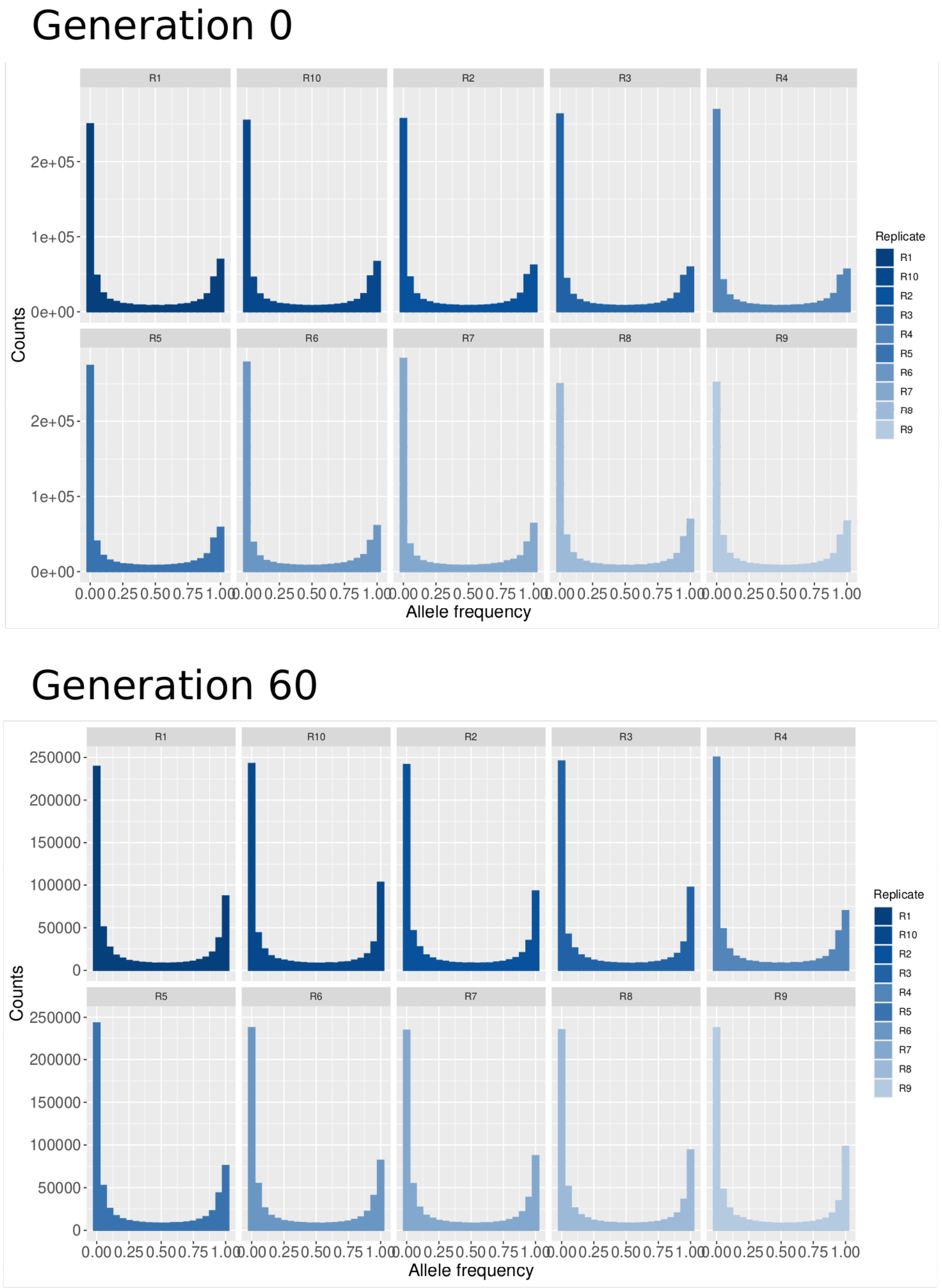


Figure S26: **Allele frequency spectra in replicate experiment #98 of the Vlachos et al. (2019) study at generations 0 and (top) and 60 (bottom), i.e., time points 1 and 7.** Each replicate population is represented in its own individual histogram plot.
